# Dynamic Single Cell Transcriptomics Defines Kidney FGF23/KL Bioactivity and Novel Segment-Specific Inflammatory Targets

**DOI:** 10.1101/2024.05.24.595014

**Authors:** Rafiou Agoro, Jered Myslinski, Yamil G. Marambio, Danielle Janosevic, Kayleigh N. Jennings, Sheng Liu, Lainey M. Hibbard, Fang Fang, Pu Ni, Megan L. Noonan, Emmanuel Solis, Xiaona Chu, Yue Wang, Pierre C. Dagher, Yunlong Liu, Jun Wan, Takashi Hato, Kenneth E. White

**Affiliations:** Department of Medical and Molecular Genetics, Indiana University School of Medicine, Indianapolis, IN, USA, 46202; Current address: The Jackson Laboratory, Bar Harbor, ME 04609, USA; Department of Medicine/Division of Nephrology, Indiana University School of Medicine, Indianapolis, IN, USA, 46202; Richard L. Roudebush Veterans Affairs Medical Center, Indianapolis, IN, USA, 46202

## Abstract

FGF23 via its coreceptor αKlotho (KL) provides critical control of phosphate metabolism, which is altered in rare and very common syndromes, however the spatial-temporal mechanisms dictating renal FGF23 functions remain poorly understood. Thus, developing approaches to modify specific FGF23-dictated pathways has proven problematic. Herein, wild type mice were injected with rFGF23 for 1, 4 and 12h and renal FGF23 bioactivity was determined at single cell resolution. Computational analysis identified distinct epithelial, endothelial, stromal, and immune cell clusters, with differential expressional analysis uniquely tracking FGF23 bioactivity at each time point. FGF23 actions were sex independent but critically relied upon constitutive KL expression mapped within proximal tubule (S1-S3) and distal tubule (DCT/CNT) cell sub-populations. Temporal KL-dependent FGF23 responses drove unique and transient cellular identities, including genes in key MAPK- and vitamin D-metabolic pathways via early- (AP-1-related) and late-phase (EIF2 signaling) transcriptional regulons. Combining ATACseq/RNAseq data from a cell line stably expressing KL with the *in vivo* scRNAseq pinpointed genomic accessibility changes in MAPK-dependent genes, including the identification of FGF23-dependent EGR1 distal enhancers. Finally, we isolated unexpected crosstalk between FGF23-mediated MAPK signaling and pro-inflammatory TNF receptor activation via NF-κB, which blocked FGF23 bioactivity *in vitro* and *in vivo*. Collectively, our findings have uncovered novel pathways at the single cell level that likely influence FGF23-dependent disease mechanisms.

**Translational statement:** Inflammation and elevated FGF23 in chronic kidney disease (CKD) are both associated with poor patient outcomes and mortality. However, the links between these manifestations and the effects of inflammation on FGF23-mediated mineral metabolism within specific nephron segments remain unclear. Herein, we isolated an inflammatory pathway driven by TNF/NF-κB associated with regulating FGF23 bioactivity. The findings from this study could be important in designing future therapeutic approaches for chronic mineral diseases, including potential combination therapies or early intervention strategies. We also suggest that further studies could explore these pathways at the single cell level in CKD models, as well as test translation of our findings to interactions of chronic inflammation and elevated FGF23 in human CKD kidney datasets.

## Introduction

Human and mouse kidneys are formed with approximately one million and 14,000 nephrons, respectively^1,2^, which are individually composed of distinct functional segments. The overall architecture and microstructure of the kidney is designed to effectively control the transport of proteins, solutes, and water between filtrate, plasma and urine^3^, thus playing a unique role in systemic mineral homeostasis. Minute to minute systemic phosphorus handling is regulated by FGF23, a hormone produced by osteocytes in response to elevated circulating phosphate or 1,25(OH)_2_ vitamin D (1,25D) (reviewed in^4^). FGF23 acts through high affinity binding to its co-receptor αKlotho (KL) and FGF receptors (FGFRs) to mediate its renal effects^5^. Under conditions of elevated blood phosphate, bone FGF23 is induced to promote phosphaturia through the downregulation of the type II sodium-dependent phosphate co-transporters Slc34a1 (Npt2a) and Slc34a3 (Npt2c) via their internalization in renal proximal tubular cells^6,7^. Additionally, FGF23 controls 1,25D in a coordinated regulation of metabolizing enzymes by decreasing the expression of the renal 1,25D anabolic *Cyp27b1* (vitamin D 1alpha-hydroxylase) and by increasing the catabolic *Cyp24a1* (vitamin D 24-hydroxylase), with the net effect of these actions to reduce both serum Pi and 1,25D. However, the full understanding of the transcriptional activity upstream of these pathways have been explored only through individual candidate regulatory studies, and are thus poorly understood.

In the kidney, FGF23 signaling through FGFR-KL complexes^8^ stimulates the mitogen activated protein kinase (MAPK) cascade. Activation of the phospho-early response kinase (p-ERK1/2) induces multiple target genes including the transcription factor early growth response gene-1 (Egr1)^5,9^. Elegant genetic mouse models have been generated through conditional deletion of KL expression from nephron proximal tubules (PT) and distal tubules (DT)^10,11^ to uncover the segment-specific roles of FGF23, FGFR and KL. However phenotypic differences across conditional mouse models have been observed, potentially due to the variability in Cre-mediated recombination efficiency and cell-type specificity of Cre expression. Overall, KL deletion in PT or DT in some models resulted in normophosphatemia or in only moderate hyperphosphatemia, compared to the severe hyperphosphatemic phenotype observed in global KL knock out mice^12,13^. Further, the molecular mechanisms responsible the altered phosphate and 1,25D regulation in diseases associated with FGF23 overexpression, such as ADHR^14^ and XLH^15^, as well as in multi-factorial disorders such as chronic kidney disease (CKD)^16^, where patients have markedly elevated FGF23 and reduced KL expression, remain incompletely understood. Finally, FGF23 signaling may be influenced by disease states, as interactions between inflammation and FGF23 have been proposed. Indeed, in a large cohort study of adults, higher quartiles of FGF23 were associated with elevated mean concentrations of IL-6, IL-10^17^. In macrophages, it was reported that FGF23 stimulated TNFα production via activation of FGFR1 signaling^18^. However, how inflammatory signals influence FGF23 bioactivity specifically in kidney cells are not well understood.

In the present study, we aimed to test the molecular function of FGF23 in kidney, the localization of these actions with regard to KL expression, and project translational manifestations to diseases associated with FGF23. To this end, renal cell identities were captured using single cell RNA sequencing (scRNAseq) on kidneys isolated from mice injected with FGF23 over multiple time points. Our line of study identified spatial and temporal transcriptional reprogramming that occurred within the nephron during FGF23-mediated signaling. We also identified *in vivo* and confirmed *in vitro* novel crossover activity of FGF23 with TNF family members and NF-κB signaling within a KL-dependent pathway. Collectively, this work isolated kidney FGF23 function at single cell resolution and identified novel pathways, genes, and transcription factors regulated by this hormone, with potential therapeutic implications for acute and chronic kidney diseases.

## Material and Methods

### Research animals

#### Ethics statement

The animal studies were reviewed and approved by Indiana University School of Medicine Institutional Animal Care and Use Committee (IACUC).

#### FGF23 administration

C57BL/6J male and female mice were purchased from the Jackson Laboratory at 9-weeks of age and housed in the IUSM Laboratory Animal Resource Center (LARC) for 1 week prior to the start of the experiments. At 10-weeks of age, mice were treated with a single intraperitoneal injection of 500 ng/g body weight of rhFGF23 (2604-FG; R&D Systems, Inc.; Minneapolis, MN) and euthanized at baseline (0h), or after 1, 4, 12, and 24h. For scRNAseq experiments, male-female kidney pairs were digested, and cells were isolated as previously described^19^. To verify the reproducibility of our approach, a duplicate of scRNAseq experiments was performed at each time point. The dataset presented in the manuscript are from a first set of experiments where a pooled of male and female kidney cells were analyzed using scRNAseq. The confirmatory analyses were tested in total kidney RNA using n=5-6 mice per time point and per gender.

#### BMS and FGF23 administration

Wild type C57BL/6 10-week-old female mice were treated with the NF-κB inhibitor BMS-345541 (0.5 mg/mouse i.p. injection, one time; S8044; BMS; Selleck Chemicals; Houston, TX), and after 1h the mice received a second single i.p. injection of FGF23; mice were then euthanized after 1h. The combined treatment duration of BMS-345541 to the mice was 2h. In another experiment, *Hyp* mice, a model of FGF23 over expression, as well as their littermates (between 8-12 weeks) were treated with a single dose of BMS-345541 (0.5 mg/mouse, by one i.p. injection) for 3h.

#### TWEAK and FGF23 administration

Wild type C57BL/6 10-week-old female mice were treated daily with recombinant tumor necrosis factor-like weak inducer of apoptosis (TWEAK 1237-TW; R&D Systems; 2 ug per mouse) for 4 days followed by a single i.p. injection of FGF23. The mice were euthanized 4 h later.

### Single cell library preparation and data processing

For each sequencing set, 10,000 cell recovery was targeted, and a single cell master mix with lysis buffer and reverse transcription reagents applied according to the Chromium Single Cell 3’ Reagent Kits V3 User Guide, CG000183 Rev A (10X Genomics, Inc.). This was followed by cDNA synthesis and library preparation. All libraries were sequenced on an Illumina NovaSeq6000 platform in paired-end mode (28bp + 91bp). The data were processed as previously described^19^. In brief, CellRanger 3.1.0 (http://support.10xgenomics.com/) was utilized to process the raw sequence data. The FASTQ files were then aligned to the mouse reference genome mm10 with RNAseq aligner STAR^20^. The aligned reads were traced back to individual cells and the gene expression level of individual genes were quantified based on the number of UMIs (unique molecular indices) detected in each cell. The filtered gene-cell barcode matrices generated by CellRanger were used for further analysis. Cells with unique gene counts over 8000 were filtered out. Gene expression was normalized by total counts from each cell, multiplied by a scaling factor 10000, and log2 transformed. The first 30 principal components from principal component analysis (PCA) were used to cluster cells by a shared nearest neighbor (SNN) modularity optimization based clustering algorithm^21^. UMAP visualization of the resulting clusters and selected genes was performed with Seurat package^22^.

### Sex hashing

It has been reported that renal cell clusters were evenly represented in kidney samples by sex except the PT clusters which exhibited gender-specific clustering^23,24^. Using this approach, the female X chromosome *Xist* gene and male Y chromosome *Kdm5d* and *Eif2s3y* genes were used as markers to distinguish male and female cells within the pooled dataset. In addition to these parameters, additional filters were applied to separate male and female cells in PT^23^. Specifically, in PT-S1, male cells were identified with markers *Slc22a28^high^*, *Cyp2e1^-^*and *Xist^-^* whereas female cells were *Spp2^high^*, *Cyp2d26 ^high^*, *Cyp4a14^-^*and *Xist^+^*. Male PT-S2 cells were identified with *Cyp2e1 ^high^*, *Nat8 ^high^* and *Xist^-^*whereas female S2 cells were predominantly *Cyp4a14 ^high^* and *Xist^+^*. The cells from PT-S3 were identified as male with *Cyp7b1^+^*, *Slc7a13 ^high^* and *Xist^-^* whereas *Slc7a12^+^* and *Xist^+^* were identified as female. Analysis of sex dependent FGF23 bioactivity was performed using a false discovery rate (FDR) less than 5%.

### Single nuclei Assay for Transposase-Accessible Chromatin data analysis

A publicly available kidney dataset, accessible in the GEO database under the number GSE210938 (GSM443123)^25^ was analyzed. Single nuclei isolation was performed following a published protocol^26^. In brief, male C57BL/6J mouse kidney was minced and homogenized in Nuclei EZ lysis buffer following a filtration through a 40-µm cell strainer and centrifuged for 5 min at 500 g. Nuclei partitioning, library preparation, and sequencing were performed following the 10X Chromium protocol. Data processing was undertaken using the Cell Ranger v6.1.2 ATAC pipeline and Loupe Browser v5.0. Fragments and peaks of snATACseq data were downloaded from GEO entry GSM6443123. Signac^27^ was employed to analyze the data using standard processes. Cells were annotated by label transfer from single cell RNAseq and Visualized using UMAP.

### Pseudotemporal ordering of single cells

To perform the pseudotime analysis on the integrated Seurat object, cells were divided into individual gene expression data files organized by previously defined cell type. R package Monocle v3 was used for dataset analysis. Outputs were obtained detailing the pseudotime cell distributions for each cell type. Positional information for the monocle plot was used to subset and color cells for downstream analyses^28^.

### Gene regulatory network inference

To identify TFs and characterize cell states, cis-regulatory analysis was employed using the R package SCENIC v1.2.4, which infers the gene regulatory network based on co-expression and DNA motif analysis. The network activity is then analyzed in each cell to identify recurrent cellular states. In short, TFs were identified using GENIE3 and compiled into modules (regulons), which were subsequently subjected to cis-regulatory motif analysis using RcisTarget with 10 kb around the TSS gene-motif rankings. Regulon activity in every cell was then scored using AUCell.

### RNA isolation and qPCR

Kidneys were harvested and homogenized in 1 ml of TRIzol reagent (15596018; Thermo Fisher Scientific; Waltham, MA) according to the manufacturer’s protocol using a Bullet Blender (Next Advance, Inc.), then further purified using the RNeasy Kit (74104; Qiagen, Germantown, MD). RNA samples were tested with intron-spanning primers specific for mRNAs expression of gene listed in Supplementary Table S1. Mouse *β-actin* or *Gapdh* was used as an internal control by RT-qPCR. The qPCR primers were purchased as pre-optimized reagents (Applied Biosystems/Life Technologies, Inc.; Waltham, MA) and the TaqMan One-Step RT-PCR kit was used to perform qPCR. PCR conditions for all experiments were 30 minutes 48°C, 10 minutes 95°C, followed by 40 cycles of 15 seconds 95°C and 1 minute 60°C. The data were collected and analyzed by a StepOne Plus system (Applied Biosystems/Life Technologies, Inc.). The expression levels of mRNAs were calculated relative to appropriate controls, and the 2^-ΔΔCT^ method described by Livak was used to analyze the data^29^.

### In vitro studies

HEK293 cells and HEK293 stably transfected with membrane bound KL (HEK293-mKL cells) were cultured in Alpha MEM (SH30265.01; Hyclone, Marlborough, MA) supplemented with 10% fetal bovine serum (S11195; R&D Systems), 1% L-Glutamine 100X Solution (Hyclone), 50,000U Penicillin-Streptomycin Solution (SV30010; Hyclone) at 37°C under 5% CO_2_ conditions. The in vitro experimental conditions were performed as follows: 2.5 x 10^5^ cells were seeded on 24-well plates (Midwest Scientific) overnight and then treated with PBS or recombinant human FGF23 for times and doses described in Results and Figure legends. In some experiments, cells were pretreated with BMS-345541 (BMS; Selleck Chemicals, Houston, TX), U0126 (MAPK inhibitor), rTWEAK (1090-TW; R&D Systems), rTNFα (210-TA; R&D Systems) prior to FGF23 (2604-FG; R&D Systems) treatment. Protein cell lysates were collected with 200 µl 1X cell lysis buffer (9803S, Cell Signaling Technology, Danvers, MA) with 0.1mM 4-(2-Aminoethyl) benzenesulfonyl fluoride hydrochloride (AEBSF, SBR00015, Sigma-Aldrich, Saint Louis, MO). Cell RNAs were extracted using ISOLATE II RNA mini kit (BIO-52073; Meridian Bioscience, Cincinnati, OH) followed by qPCR analysis as described above.

#### Bulk mRNA sequencing

Total RNA from HEK293-mKL cells treated with vehicle or FGF23 (50 ng/mL for 4 h) was extracted and evaluated for its quantity and quality using an Agilent Bioanalyzer 2100; 100 ng of total RNA was used for the cDNA libraries. Library preparation included mRNA purification/enrichment, RNA fragmentation, cDNA synthesis, ligation of index adaptors, and amplification, following the KAPA mRNA Hyper Prep Kit Technical Data Sheet, KR1352 – v4.17 (Roche Corporate). Each resulting indexed library was quantified, and its quality accessed by Qubit and Agilent Bioanalyzers; multiple libraries were pooled in equal molarity. The pooled libraries were denatured and neutralized before loading on a NovaSeq 6000 sequencer at 300 pM final concentration for 100b paired-end sequencing (Illumina, Inc.). Approximately 30-40M reads per library were generated. A Phred quality score (Q score) was used to measure the quality of sequencing. More than 90% of the sequencing reads reached Q30 (99.9% base call accuracy). The sequencing data were first assessed using FastQC (Babraham Bioinformatics, Cambridge, UK) for quality control. The sequencing reads were mapped to the human genome hg38 using STAR (v2.7.2a) with the following parameter: ‘--outSAMmapqUnique 60’ ^20^. Uniquely mapped sequencing reads were assigned to Gencode M22 gene using featureCounts (v1.6.2) ^30^ with the following parameters: “–p –Q 10 –O”. The genes were kept for further analysis if their read counts > 10 in at least 3 of the samples, followed by the normalization using TMM (trimmed mean of M values) method and subjected to differential expression analysis using edgeR (v3.24.3) ^31^. Gene Ontology and KEGG pathway functional enrichment analysis was performed on selected gene sets, e.g., genes undergoing both significant differential expressions and notable changes of open chromatin accessibilities, with the cut-off of false discovery rate (FDR) < 0.05 using DAVID ^32^. Canonical pathways from RNAseq data were generated through the use of IPA (QIAGEN Inc., https://www.qiagenbioinformatics.com/products/ingenuity-pathway-analysis)^33^; n=4 samples per condition.

#### Bulk Assay for Transposase-Accessible Chromatin sequencing (ATACseq)

After treatment with FGF23 or vehicle for 4 h (see Results) cells were washed twice in 1X PBS, then dissociated with trypsin (SH30042, Hyclone) for 5 minutes. Cells were resuspended in ice cold 1X PBS, dead cells were removed, and live cells processed for nuclei isolation and ATAC sequencing according to published protocols ^34^. Briefly, cells were collected in cold PBS and cell membranes were disrupted in cold lysis buffer (10 mM Tris–HCl, pH 7.4, 10 mM NaCl, 3 mM MgCl2 and 0.1% IGEPAL CA-630). The nuclei were pelleted and resuspended in Tn5 enzyme and transposase buffer (Illumina Nextera® DNA library preparation kit, FC-121-1030). The Nextera libraries were amplified using the Nextera® PCR master mix and KAPA biosystems HiFi hotstart ready-mix successively. AMPure XP beads (Beckman Coulter) were used to purify the transposed DNA and the amplified PCR products. The resulting ATACseq libraries were sequenced on Illumina NovaSeq 6000 and paired-end 50 bp reads were generated. Illumina adapter sequences and low-quality base calls were trimmed off the paired-end reads with Trim Galore v0.4.3. Bowtie2 ^35^ was used for ATACseq read alignments on the human genome (hg38). Duplicated reads were removed using Picard developed by the Broad Institute [https://broadinstitute.github.io/picard/ (Accessed: 2018/02/21; version 2.17.8)]. Low mapping quality reads and mitochondrial reads were discarded in further analysis. Peak calling of mapped ATACseq reads were performed by MACS2 ^36^ with a Bonferroni adjusted cutoff of p-value less than 0.01. Peaks called from multiple samples were merged, after removing peaks overlapping with ENCODE blacklist regions ^37,38^. Reads locating within merged regions in different samples were counted by pyDNase ^39^. The data was filtered using at least 10 cut counts in more than one of the samples, then normalized using TMM (trimmed mean of M values) method and subjected to differential analysis using edgeR (v3.24.3) ^31,40^. Motif enrichment of differential accessibility peaks with a false discovery rate cut-off of 0.05 was performed using Homer ^41^.

### Immunoblotting

HEK293-mKL cells were lysed with 300 µL 1X Lysis buffer (Cell Signaling Technology, Inc., Danvers, MA, USA; 9803-S) with 1 µg/mL 4-(2-aminoethyl) benzenesulfonyl fluoride hydrochloride protease inhibitor (AEBSF; SBR00015, Sigma-Aldrich, Inc.;). Total cell lysate protein concentrations were determined with the Better Bradford Kit (23236; Thermo-Fisher Scientific) according to the manufacturer’s instructions. Western blot analysis was performed with 30 µg of cellular lysates. The blots were incubated with 1:1000 primary antibody to pERK (T202/Y204; 4370S; Cell Signaling Technologies, Danvers, MA, USA) tERK (9102S; Cell Signaling Technologies, Danvers, MA, USA), p-NF-κB-p65 (S536; 93H1; Cell Signaling Technologies, Danvers, MA, USA), NF-κB-p65 (D14E12; Cell Signaling Technologies, Danvers, MA, USA), p-c-JUN (S73; D47G9; Cell Signaling Technologies, Danvers, MA, USA), t-c-JUN (90A8; Cell Signaling Technologies, Danvers, MA, USA), p-c-FOS (Ser32; D82C12; Cell Signaling Technologies, Danvers, MA, USA), and c-FOS (9F6; Cell Signaling Technologies, Danvers, MA, USA) overnight then incubated with secondary antibody at 1:2000 (anti-rabbit– horseradish peroxidase [HRP]; Cell Signaling Technologies). Blots were stripped using SDS-glycine and reprobed with 1:10,000 anti-β-actin-HRP (A3854; Sigma-Aldrich). Detection was performed using the ECL Prime Western Blotting Detection Reagents (Amersham-GE Healthcare, Pittsburgh, PA, USA) and a GE AB1600 digital imager.

### Quantification and statistical analysis

As described in Results and Methods, the most recent updated R software packages with robust affiliated statistics were used to analyze the scRNAseq and snATACseq datasets. Statistical analyses of the *in vitro* data were performed by one-way ANOVA to assess the differences between the same cell line responses to FGF23 treatment and a two-way ANOVA to assess the response differences between two different cell lines. Statistical analyses of the *in vivo* data presented were performed by one-way ANOVA to assess the differences between the same gender in response to treatment and a two-way ANOVA to assess the differences between genders and treatments. Means and standard deviations were used in bar graphs and kinetic curves. Significant changes were considered when *P* <0.05.

## Results

### Renal scRNAseq enables sex-specific sample demultiplexing

FGF23-mediated control of renal phosphate and vitamin D metabolism is essential for maintaining, in the short term, proper systemic energetics through ATP production, as well as over the long term, a mineralized skeleton. With the goal of testing dynamic FGF23 bioactivity within the kidney nephron, male and female C57BL/6 mice were injected with recombinant FGF23 (400 ng/g) for 0, 1, 4, and 12h followed by euthanasia, and the generation of a pooled male/female scRNAseq dataset (Fig. 1a). We leveraged the currently available computational methodology to map sexually dimorphic kidney phenotypes through counting the number of sequencing reads of male-*versus* female-specific genes^23^. It has been reported that renal cell clusters were evenly represented in kidney samples by sex except PT cell populations which we confirmed exhibit gender-specific clustering^23,24^. Male and female cell types were demultiplexed within clusters using the sex-specific markers, and analyzed together as there were no sex-dependent transcriptional changes in response to FGF23 (see Methods). By applying dimensionality reduction *via* Uniform Manifold Approximation and Projection (UMAP), 21 distinct clusters were identified in kidney including epithelial, endothelial and immune cell clusters, which were classified in unique populations based upon the known expressional markers of specific renal cell types (Fig. 1b-d)^19,23,42–45^. To assess the effect of acute FGF23 administration on renal cell populations, stacked bar plots were used to visualize the proportion of different cell types. Proximal tubule S1-S3 cells represented the most abundant types (Fig. 1e), consistent with this segment comprising the predominant kidney segment. FGF23 treatment did not affect the overall epithelial cell population distributions (Fig. 1d).

**Fig. 1:**
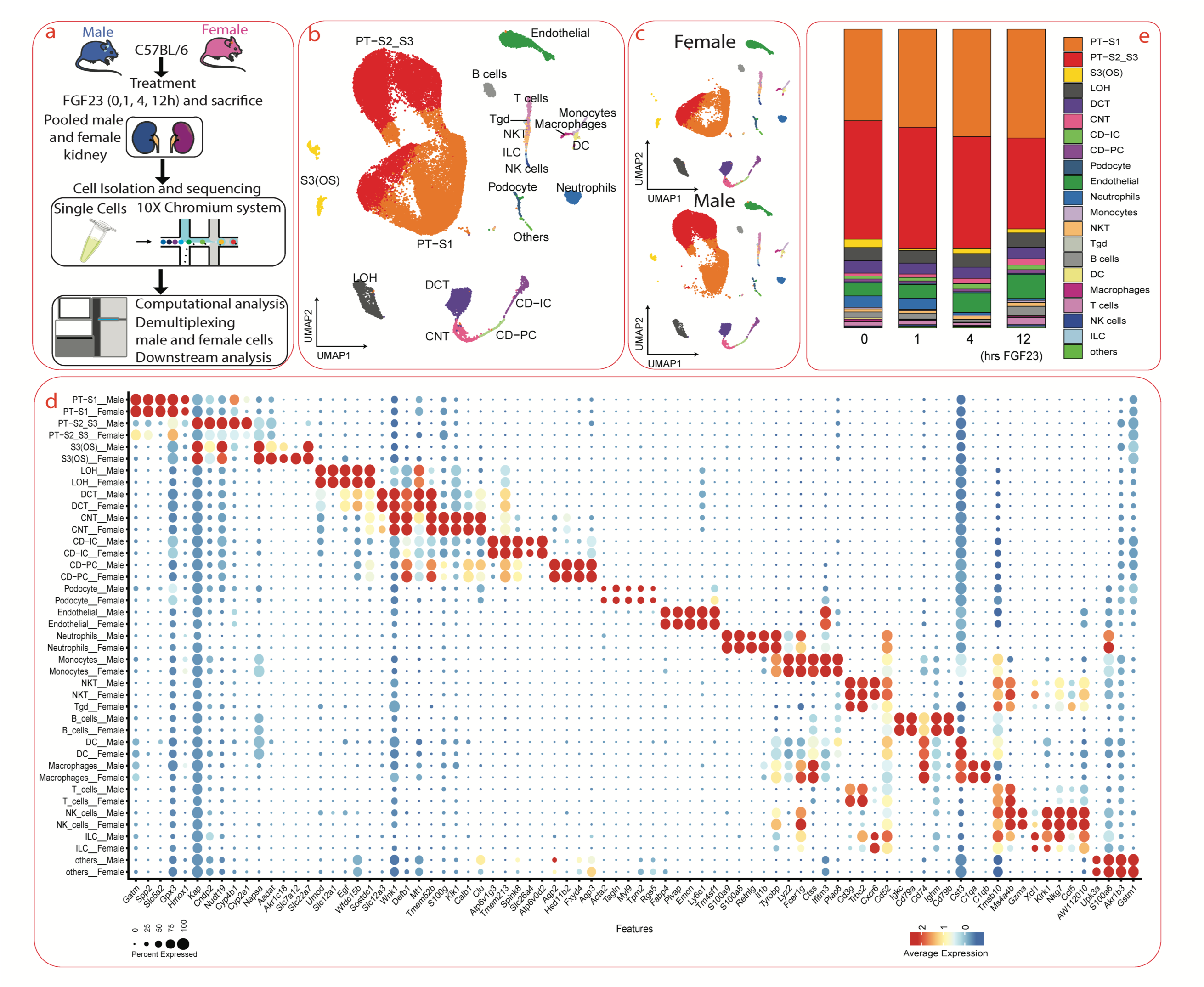
Renal sex-multiplexing scRNA sequencing. **a.** Experimental Overview. Pooled male and female kidneys from FGF23-injected mice were dissociated into single-cell suspensions followed by scRNAseq and downstream computational and molecular analysis. **b.** Unsupervised UMAP clustering identified various renal cell populations divided into 21 distinct cell types. **c.** Known sex-specific markers (described in Results) were used to identify male and female cell populations. **d.** Dot plot of representative genes show sex-specific cell types. **e.** Stacked bar plot displays the relative proportions of each cell-type after FGF23 treatment. The different cell types identified are color-coded and annotated for clustering as in Fig. 1b.

### Klotho expression coordinated with FGF23 activity in the nephron

The full extent of FGF23 bioactivity within the nephron remains unresolved as its co-receptor Klotho (KL)^5^ is expressed in multiple segments. FGF23 is known to elicit potent bioactivity to reduce both renal phosphate reabsorption and production of active 1,25D in proximal tubule (PT), formed by segments S1, S2, and S3. KL is highly expressed in the distal tubule (DT) which is composed of the distal convoluted tubule (DCT) and connecting tubule (CNT; reviewed in^4^), however cell type-specific actions of KL remain unresolved. In testing *KL* expression across the nephron, *KL* was undetectable in podocytes. In PT cells, *KL^high^* cells were predominant in PT-(S1, S2, S3) and the proportion of *KL^high^* cells fell during transition to the loop of Henle. The highest expression of *KL* was detected in DCT and CNT, in agreement with previous localization approaches using immunohistochemistry and immunofluorescence^9,11,46,47^. The collecting duct intercalated cells were predominantly *KL^low^* cells, as were the immune and endothelial cells identified in the dataset (Fig. 2a and Supplementary Fig. 1). There was no significant difference for *KL* expression in renal cells when comparing male *vs* female cells across epithelial cells (Supplementary Fig.1). The expression of *Fgfr1* was widely detected in renal epithelial cells (Supplementary Fig. 2), confirming the ability of these cells to potentially form FGF23-FGFR1-KL complexes, known to trigger robust FGF23 intracellular signaling. Of note, previous studies demonstrated that deletion of *Fgfr1* from PT and DT cells disrupted renal phosphate and calcium transport^48^.

**Fig. 2:**
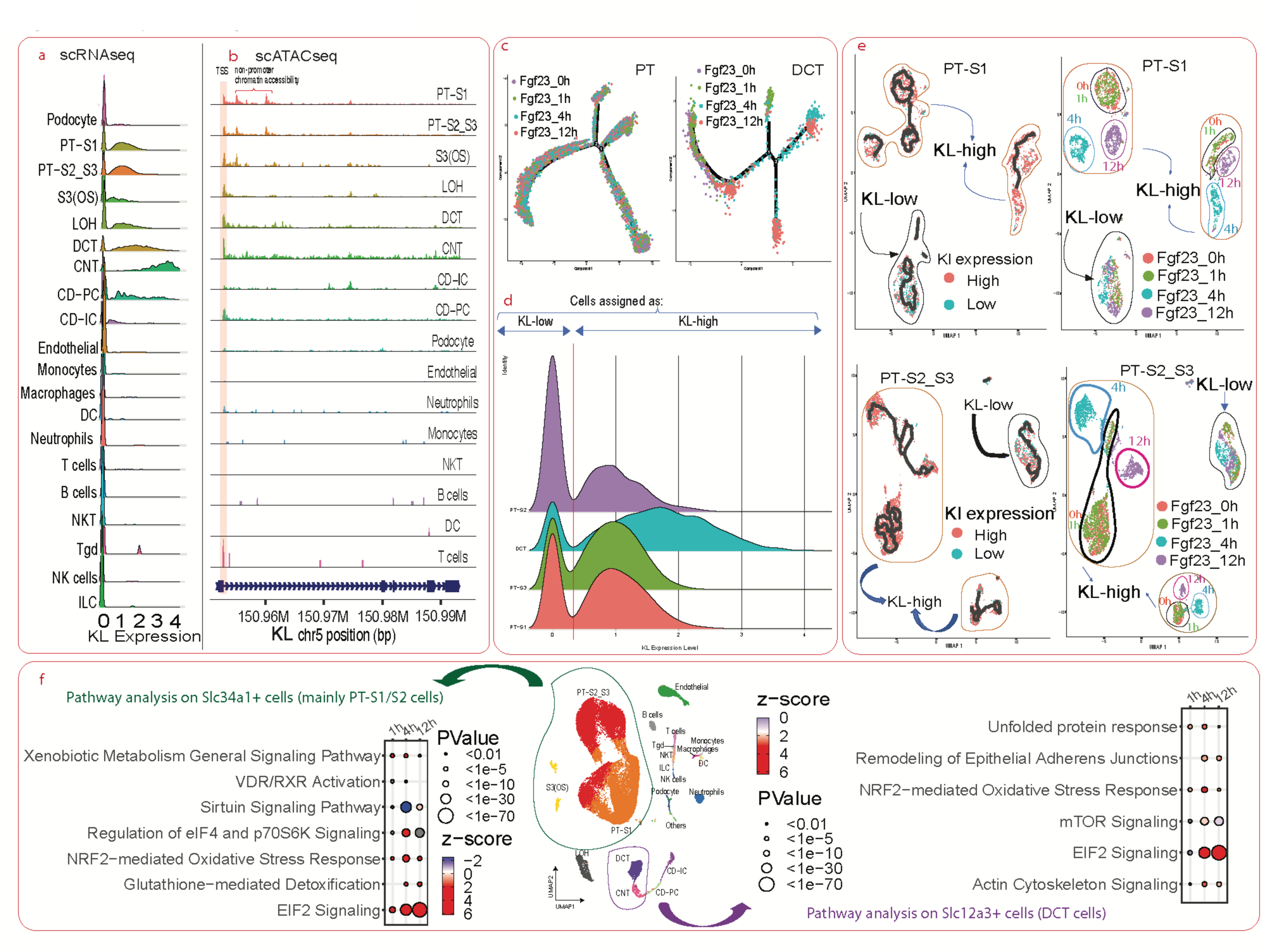
KL defines a specific FGF23 signature in renal tubules. **a.** Ridgeplots show *KL* mRNA expression in select renal cells. **b.** snATACseq detected a differential chromatin accessibility between PT, LOH, DCT, and CNT *versus* Stromal and Endothelial cells at the KL promoter region. The red bracket shows differential chromatin accessibility in a region located between exon 1 and exon 2 when comparing PT cells types *versus* LOH, DCT, CNT, and CD-IC. **c.** Cell states in PT and DCT cells in response to FGF23 with a time-dependent clustering in DCT versus PT. **d.** Ridgeplot analysis shows KL expression in PT(S1-S3) and DCT. The vertical red line demarcates cells with low KL expression (KL^low^) or high KL expression (KL^high^). **e.** The Monocle program was used to cluster PT-S1 and PT-S2_S3, to separate cells into KL^high^ and KL^low^ groups, and display cells by treatment condition (0, 1, 4, and 12h). **f.** Overview of nephron segment specific pathways activated in proximal tubule and distal tubule cells in response to FGF23. Ingenuity Pathway Analysis (IPA) was used to predict the statistically significant canonical pathways upregulated in PT and DT in response to FGF23 at 1, 4, and 12h. The z-score represents the ratio of the number of upregulated genes found in each pathway over the total number of genes known to be involved in that pathway; the size of the dot represents the p value.

Since KL expression was restricted to epithelial cells (Fig. 2a and Supplementary Fig. 1), next it was sought to determine whether cell-specific predisposition mechanisms were present at the genomic level to potentially regulate KL expression. We performed analysis of single nucleus assay for transposase-accessible chromatin sequencing (snATACseq) of mouse kidney which identified the expected cell clusters including epithelial cells (Supplementary Fig. 3a). We found that the accessible KL promoter regions were limited to PT, loop of Henle (LOH), DCT and CNT cells, supporting our findings of *KL* mRNA distribution within the nephron (Fig. 2a-b). Interestingly, it was observed that within the *KL* gene body, differentially accessible chromatin regions between exons 1 and 2 were present. At these regions, the chromatin profile was accessible in PT but closed in LOH, DCT, and CNT cells (Fig. 2b and Supplementary Fig. 3b-d). Furthermore, the differential chromatin accessibility at the site nearer to the *KL* promoter corresponded to a VDR binding site (Supplementary Fig. 3c). Taken together, the differential genomic predisposition across renal cells within the *KL* gene supports FGF23 site-specific regulation of mineral metabolism.

Next, we applied pseudotime analysis^28^ to the integrated 4 datasets (Fgf23 delivery at 0h, 1h, 4h, and 12h). We identified a more heterogenous cell-state in PT regardless of FGF23 treatments and more homogeneous cell-populations (DCT) that clustered depending upon FGF23 treatment (Fig. 2c). In DCT, a more profound phenotype was observed at the 4h time point when compared with untreated cells. At the 12h treatment time point, although most DCT cells were still clustered into a distinct branch (Fig. 2c, red cells), some cells returned to the original branch corresponding to untreated samples (Fig. 2c). It was then hypothesized that the FGF23-dependent cell-state clustering was driven by differential KL expression. To map the distribution of KL across the kidney, PT and DT cells were used as reference controls. Interestingly, using the visualization tool Ridgeplot, *KL* mRNA on the PT and DCT cell subpopulations showed a clear cutoff if *KL* expression was >0.4 or <0.4 (Supplementary Fig. 1a), allowing the assignment of renal cells as *KL^high^* or *KL^low^*, respectively. This approach provided the opportunity to refine the identification of cell specific KL-dependent FGF23 signaling pathways with higher confidence. Cells were then assigned to KL^low^ or KL^high^ to determine cellular states in response to FGF23 (Fig. 2d) and further used Monocle 3 to discover subtypes of cells that clustered based upon KL expression. To build upon the results above and determine the role of KL-dependent FGF23 bioactivity on cell phenotype changes in the PT, we circumvented the confounding presence of the heterogenous PT cell population for KL expression levels by parsing PT-S1 and PT-S2/S3 cells into *KL^high^* and *KL^low^* subclusters (Fig. 2e top and bottom left) followed by testing FGF23-mediated effects between cell populations. Interestingly, *KL^high^* subclusters were driven by clear spatio-temporal separation of cells in response to FGF23. These data showed that when stimulated by FGF23, the *KL^high^* cells possessed distinct transient cellular phenotype changes in contrast to *KL^low^*clusters, suggesting differential gene signatures driven by FGF23 in presence or absence of KL (Fig. 2e top and bottom right). In contrast to PT-S1 or PT-S2_S3 cells, Monocle analysis on DCT cells revealed only a single partition, consistent with the homogeneity of the DCT cell population, as almost all cells expressed KL. Further, in DCT cells, the time-dependent status of FGF23 activity was similar to KL-dependent FGF23 bioactivity in PT cells, highlighting that KL-dependent FGF23 signaling may have conserved mechanisms across nephron segments (Supplementary Fig. 4a-c). Collectively, these findings support the concept that KL production within a segment-specific cell population can be used to isolate components of downstream FGF23 bioactivity, and that individual cells that have more highly expressed KL act in *cis* as a prerequisite for FGF23 function.

The proximal tubule (PT) and distal tubule (DT) segment-specific signaling pathways associated with transcriptional changes in response to FGF23 were next tested. To identify common and unique pathways driven by FGF23 bioactivity in Slc34a1 (Npt2a) positive cells (the highest expression is observed in PT-S1) and Slc12a3 positive cells (primarily DCT), differentially upregulated genes in PT-S1 cells *vs* DCT cells were first isolated (Supplementary Figure 5a) then the Ingenuity Pathway Analysis (IPA) tool was used to identify enriched and upregulated pathways. As expected, FGF23 signaling was associated with MAPK pathways such as EIF2 (genes *Ddit3* and *Atf3/Atf4*) in both PT-S1/S2 and DCT cells (Fig. 2f). The EIF2 signaling was consistently elevated across time with FGF23 treatment. In addition to the activation of EIF2 signaling, other stress-related pathways such as NRF2-mediated oxidative stress response and the unfolded protein response were increased in both PT-S1/S2 and DCT. Besides the genes associated with stress response, known FGF23-mediated pathways such as VDR signaling was induced with FGF23 treatment in PT cells via significant upregulation of genes such as *CEBPA, CYP24A1, HES1, NCOR1, PPARD, PSMC5, and VDR* (*P* < 0.01; expression of Vdr in PT-S1 is shown in Supplementary Fig. 5b). Collectively, our temporal data support that FGF23-mediated vitamin D metabolism is preceded by a broad and overlapping biological process initiated by MAPK and EIF2 signaling (Supplementary Fig. 5c).

### Dynamic, cell specific FGF23 bioactivity

We next sought to localize FGF23 bioactivity across the nephron at 1, 4, and 12h after FGF23 administration with a specific focus on key FGF23 targets, as well as to identify novel FGF23 regulated genes. First, we assessed the influence of FGF23 treatment on KL regulation since this co-receptor is required for high affinity FGF23 bioactivity. At the mRNA level, *KL* expression remained largely stable in response to acute FGF23 administration, although a trend of modest *KL* decrease was noted at 4 and 12h after treatment (Fig. 3a). *Egr1,* a transcription factor and well-known FGF23 target^49^ acutely rose primarily in PT, DCT and CNT 1h after FGF23 delivery. These increases in *Egr1* observed at 1h decreased at 4h before resetting completely to baseline at 12h, suggesting a resolution of the 1h “*early phase*” of FGF23 actions (Fig. 3b). Only epithelial nephron cells responded to FGF23 delivery as we observed no significant increase of Egr1 in immune or endothelial cells which were *KL^low^*cell populations. In whole kidney, *Egr1* expression was confirmed as significantly elevated in response to FGF23 at 1h. In the bulk renal RNA analysis, *KL* expression was not statistically different after FGF23 treatment (Fig. 3c-d).

**Fig. 3:**
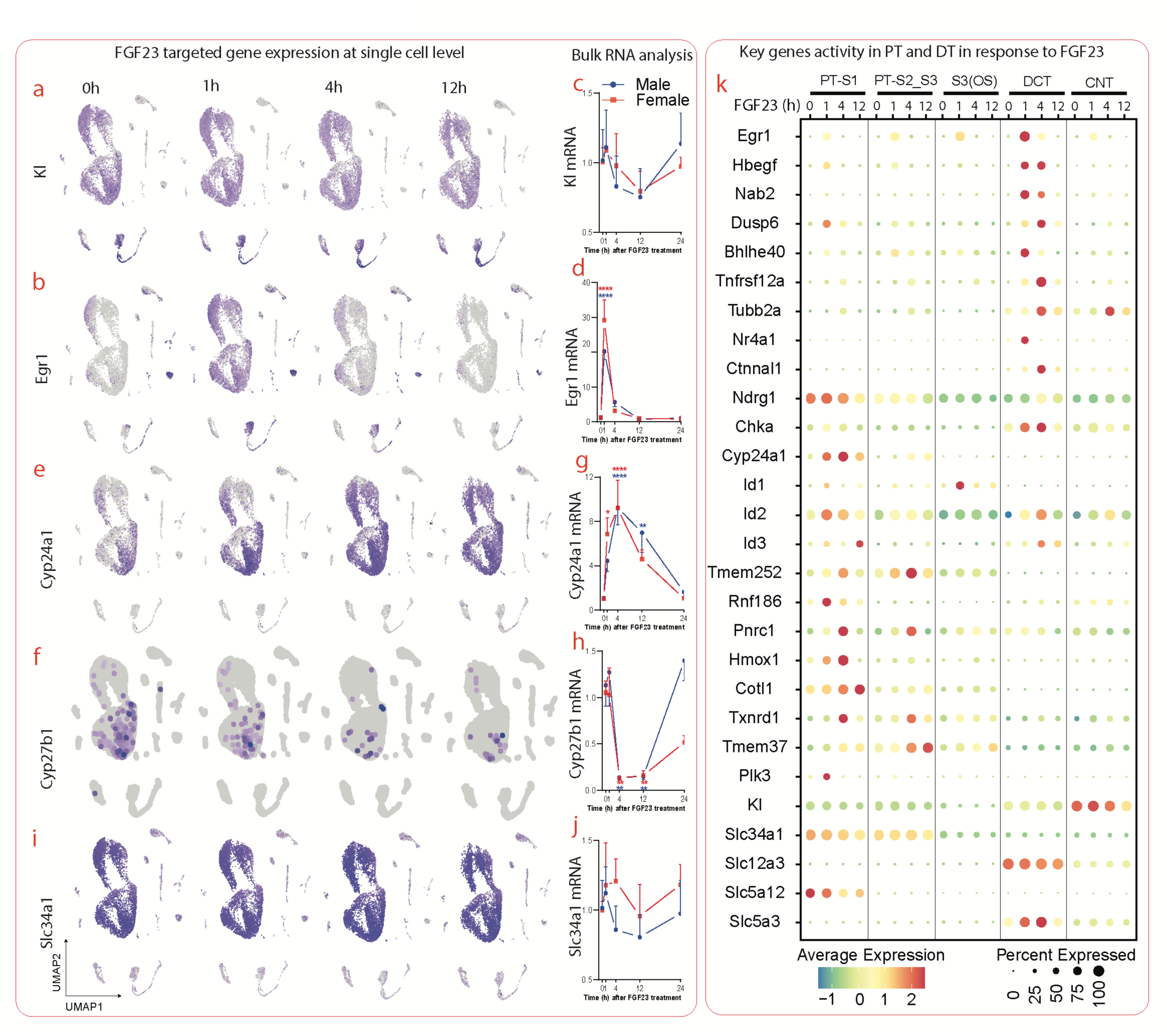
FGF23 bioactivity across the nephron. **a-j.** The mRNA expression of FGF23 target genes *Kl*, *Egr1, Cyp24a1, Cyp27b1,* and *Slc34a1,* respectively, was tested in the kidney scRNAseq dataset and by qPCR analysis. Feature plots show the expression of FGF23 targets in the different renal cell populations. The kinetic curves represent the qPCR analysis in total kidney with male data shown in blue and female data represented in red. The qPCR data are expressed as fold change (2-ΔΔCt) relative to the housekeeping gene β-actin. Data are shown as mean +/- standard deviation. **k.** Dot plot of representative genes that define FGF23 bioactivity in PT(S1-S3), DCT, and CNT.

FGF23 is known to reduce systemic concentrations of the active form of vitamin D, 1,25D, through the induction of Cyp24a1 and the repression of Cyp27b1^50–53^. To pinpoint the cells involved in the regulation of 1,25D metabolic enzymes by FGF23, *Cyp24a1* and *Cyp27b1* mRNAs were mapped in the scRNAseq dataset. The constitutive and specific expression of *Cyp24a1* and *Cyp27b1* mRNA were localized to PT segments S1, S2 and S3. In response to FGF23, *Cyp24a1* mRNA increased in PT, reaching a maximal level at 4h whereas *Cyp27b1* expression decreased (Fig. 3e and 3f). In assessing the temporal responses of FGF23 bioactivity in total kidney using qPCR analysis, *Cyp24a1* mRNA was confirmed to increase in response to FGF23 and reached a peak at 4h. At 12h after FGF23 injection, *Cyp24a1* expression was reduced compared to 4h, although these levels were still elevated in comparison to baseline (0h). At 24h after treatment, renal *Cyp24a1* mRNA returned to baseline. In contrast to *Cyp24a1*, renal *Cyp27b1* expression was dramatically suppressed between 4 and 12h before recovery at 24h (Fig. 3g and 3h). The expressional mapping of *Slc34a1* (Fig. 3i and 3j) showed little change at the transcriptional level, in accord with reports that Npt2a is largely regulated via protein withdrawal/insertion in the PT apical membrane. Indeed, we found that FGF23 treatment resulted in a time-dependent decrease of Slc34a1 protein expression over the 24h time course (Supplementary Fig. 6). Genes such as the inhibitor of DNA binding (*Id1-3*) were highly upregulated in PT whereas the *Ngfi-A* binding protein 2 (Nab2), and *Mcl1* were increased in DT (Fig. 3k). Other genes such as *Egr1*, and *Hbegf* were induced in both nephron segments as described above (Fig. 3k). Finally, a cell survival transcriptional phenotype was associated with the upregulation of genes such as *Hes1*, *Tmem208*, and *Myof*, in parallel with up-regulation of TNFR family members and NF-κB dependent genes such as *Tnfrsf12a*. Collectively, these results support that KL-dependent FGF23 signaling rapidly and transiently induced an acute phase of signaling that overlaps in PT and DT cells, as well as a more latent phase that includes site-specific, dynamic control of 1,25D metabolism.

### Identification of temporally-driven FGF23 transcriptional regulons

As described above, pathway analyses predicted differential gene regulation downstream of FGF23-FGFR1-KL interactions. To determine the renal transcription factors (TFs) activated in response to FGF23 signaling, TF regulon activity was examined using Single-Cell rEgulatory Network Inference and Clustering (SCENIC)^54,55^. Using inferred gene correlation networks followed by motif-based filtration, SCENIC selects direct targets of each TF as sets of regulons. We applied SCENIC to PT-S1 cells and identified enriched regulon states in response to FGF23 treatments at 1, 4, and 12h. The top 15 regulons at 1h were *Klf10*, *Nr1h4*, *Foxo3*, *Bhlhe40*, *Atf4*, *Stat3*, *Myc*, *Nr3c1*, *Egr1*, *Junb, Jun, E2f3, Mafg, Fos, and Jund* (Fig. 4a; and Supplementary Data 1). At 4 and 12h after FGF23 treatment, *Nfkb1* and *Nfkb2* regulons were increased (Fig. 4a; and Supplementary Data 1). *Egr1, Fos* and *Bhlhe40* regulon activity and mRNA expression were activated in specific cell subsets (Fig. 4b-d; cells surrounded in red). Similar to the regulation of PT-S1 cell *Egr1, Fos,* and *Bhlhe40* in response to FGF23 in the mice, in vitro kinetic studies confirmed that these same mRNAs were upregulated in human HEK293-mKL cells (HEK293 renal embryonic cells stably expressing KL) with FGF23 treatment (Fig. 4e-g).

**Fig. 4:**
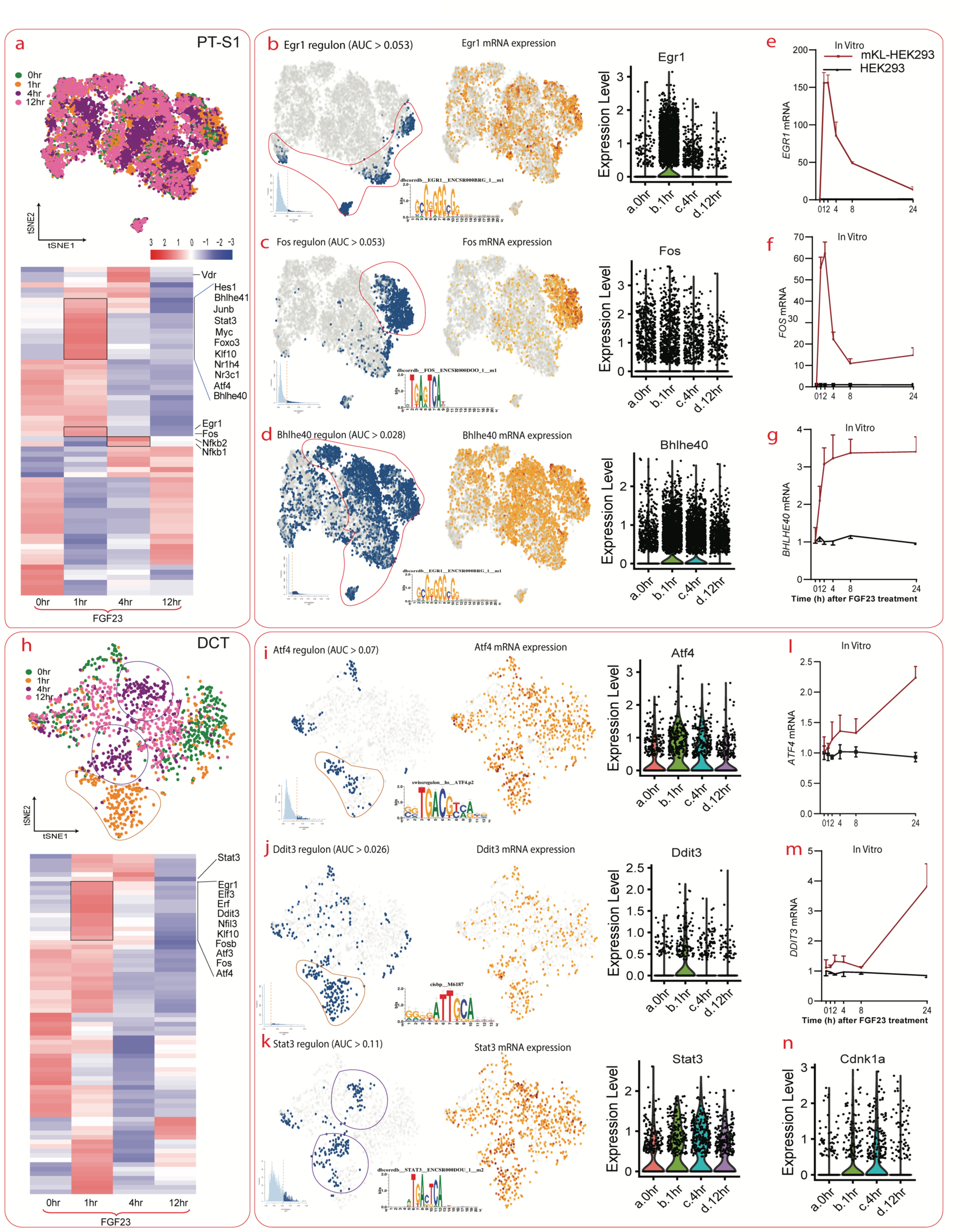
Regulon activity induction in response to FGF23. **a-Top.** SCENIC analysis was performed on PT-S1 cell data derived from FGF23-treated mice at different time points, and the cells dimensionally reduced in a t-distributed stochastic neighbor embedding (t-SNE) plot. **a-Bottom.** The heatmap shows regulon activities in PT-S1 at 0h, 1h, 4h, and 12h after FGF23 treatment. The red coloring represents the highest regulon activity. **b-d.** t-SNE plots illustrate regulon activities of *Egr1*, *Fos*, and *Bhlhe40* (blue) of PT-S1 cells in parallel with the transcription factor mRNA expression detected in the SCENIC-generated dataset (orange color in t-SNE plot). The violin plots show the corresponding mRNA expression in the original scRNAseq dataset. The insets illustrate the AUCell score distribution for a respective regulon and the predicted binding motifs. **e-f.** The mRNA expression of *EGR1, FOS,* and *BHLHE40* obtained from *in vitro* studies were measured by qPCR. The red kinetic curve represents the HEK293-mKL cell studies whereas the black kinetic curve shows the data from native HEK293 cell experiments. **h-Top.** SCENIC analysis was performed in DCT cells derived from the FGF23-treated mice at different time points. **h-Bottom.** The heatmap graphic shows regulon activities in DCT at 0h, 1h, 4h, and 12h following FGF23 administration. The red color represents the highest regulon activity. **i-k.** t-SNE plots show the regulon activity of *Atf4*, *Ddit3*, and *Stat3* in DCT in blue in parallel with the transcription factor mRNA expression (t-SNE plot with orange dots) in the SCENIC dataset. The violin plots display mRNA expression in the scRNAseq dataset. The insets illustrate the AUCell score distribution for the regulon and the predicted binding motifs. **l-m.** The kinetic curves show the mRNA expression of *ATF4,* and *DDIT3*. The red kinetic curve represents the mKL-HEK293 cell line, and the black kinetic curve highlights the HEK293 parent cells. **n.** The violin plot shows the mRNA expression of *Cdkn1a* in the scRNAseq dataset.

In the DCT cell population, overlapping regulon activity with PT cells was observed at 1h, including transient enrichments in *Egr1*, *Elf3*, *Erf*, *Ddit3*, *Nfil3*, *Klf10*, *Fosb*, *Atf3*, *Fos*, and *Atf4*. At 4 and 12h after treatment, these initial regulon increases reset to baseline (Fig. 4h; and Supplementary Data 2), and new downstream regulons including *Vezf1*, *Elk4*, and *Stat3* were induced at 4h. These data support a *‘late phase’* of FGF23-related transcription factors. Importantly the regulon activities of the transcription factors downstream EIF2 signaling, *Atf4* and *Ddit3,* were induced 1h after FGF23 treatment (Fig. 4i-j; and Supplementary Data 2), and Stat3 regulons were specifically activated in the 4h FGF23 treatment cell subset (Fig. 4k; cells surrounded in purple). *ATF4*, and *DDIT3* were confirmed to be upregulated in HEK293-mKL cells in response to FGF23 (Fig. 4l-m), and the *Stat3* downstream gene *Cdkn1a* was increased in DCT (Fig. 4n), confirming these actions. Besides the increase of *Egr1* mRNA levels in PT-S1 (Fig. 4b) and DCT (Supplementary Fig. 7a), the differential mRNA expressions of other TFs were more modest, although the regulon-based analysis showed a clear enrichment. For example, Fos regulons in PT-S1 (blue-labelled cells in Fig. 4c) and in DCT (blue-labelled cells in Supplementary Fig. 7b) were clearly enriched in the 1h-treated cells. In sum, these data demonstrate that FGF23 bioactivity coordinated an overlapping panel of early- and later-phase transcriptional activators and repressors that were specific to distinct nephron cell populations.

### FGF23-induced NF-κB is a negative regulator of MAPK signaling

Next, we focused on potential FGF23-mediated MAPK and NF-κB crossover signaling as supported by observations including: 1) an increase of NF-κB targets such as *Tnfrsf12a* in response to FGF23 in both PT and DT (Fig. 3k); 2) detection of a sustained enrichment of NF-κB TFs at 4 and 12h in response to FGF23 using SCENIC (Fig. 5e); and 3) the important translational impact of NF-κB signaling in a broad group of inflammatory renal diseases. To test the transcriptomic and genome accessibility responses to FGF23 in isolation, we performed RNAseq and ATACseq experiments using HEK293-mKL cells that were treated with either FGF23 (50 ng/ml) or vehicle (Control) for 4h (see schematic in Fig. 5a). Our analysis identified 1844 differentially expressed genes relative to the control condition (Fig. 5b), including many of the predicted ‘early-phase’ gene transcripts within the MAPK pathway. The volcano plot highlights key known and novel genes regulated by FGF23 including upregulated genes such as *EGR*1-4, *FOS*, *FOSL1, FOSL2, JUNB, HBEGF, NAB2, ATF3, MAFF, TNFRSF1A*, and *TNFRSF12A* (Fig. 5b). The primary ATACseq data analysis identified 8235 and 1334 significantly increased or decreased chromatin accessibilities, respectively, that corresponded to the identified open chromatin peaks in the dataset (Fig. 5c, volcano plot). The open chromatin map was found to be reproducible across the replicates in independent samples and libraries (see Supplementary Fig. 8a). By examining the peak changes with respect to their location relative to known annotated genomic features, 15.41% and 11.02% of these peaks were associated with increased and decreased chromatin accessibilities, respectively, and located within 10kb upstream of the transcription start site (Fig. 5c, pie chart).

**Fig. 5.**
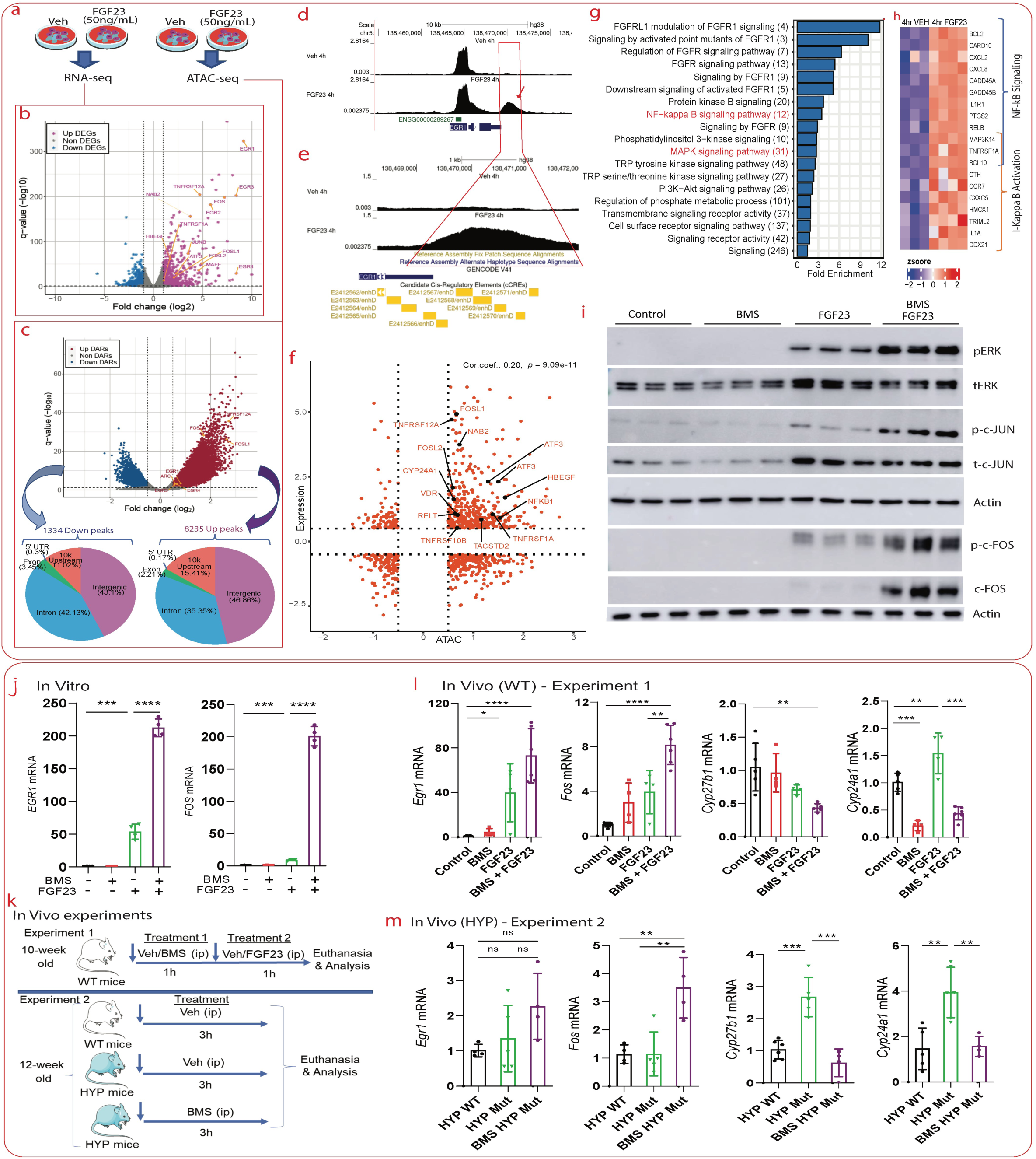
NF-kB is a negative regulator of FGF23 bioactivity. **a.** Schematic representing HEK293-mKL cell treatment with FGF23. **b-c.** Following the treatment, RNAseq and ATACseq were performed, and violin plots were generated to isolate differentially expressed genes and chromatin accessibility. The pie charts under the ATAC violin plots show the repartition of peaks along the genome. **d.** Chromatin accessibility at the EGR1 genomic region shows increased accessibility with FGF23 treatment. **e.** An enlargement at the distal EGR1 enhancer region isolated the genomic coordinates within the regions of FGF23-induced chromatin accessibility increases. **f.** Integration of ATACseq and RNAseq data displays selected genes that showed both increased chromatin accessibility (x-axis) with gene expression alterations (y-axis). **g.** Functional enrichment analysis using the Database for Annotation, Visualization and Integrated Discovery (DAVID) on upregulated DEGs with FGF23 treatment (numbers in parenthesis are gene counts). **h.** Heatmap displays the genes associated with NF-κB pathway activation. **i.** HEK293-mKL cells were pretreated with BMS-34554 (10 μM) for 1h followed by FGF23 (50 ng/ml) for 4h. Immunoblots were then performed to assess the phosphorylation of ERK, cJUN, and cFOS. **j.** HEK293-mKL cells were pretreated with BMS-34554 (10 μM) for 1h followed by FGF23 (50 ng/ml) for 4h and subsequent *EGR1* and *FOS* mRNA expression analysis by qPCR. **k.** Experimental design shows the workflow of the treatment performed by treating C57BL/6 or Hyp mice with BMS and/or FGF23. **l-m.** qPCR analysis of mouse *Egr1, Fos, Cyp27b1,* and *Cyp24a1* following treatments.

Within the EGR1 genomic region, in response to FGF23 a transient increase of chromatin accessibility at a distal enhancer region (E2412567-2412571/enhD) was present, in contrast to baseline where these regions located at the intergenic region between *EGR1* and *CTNNA1* were closed (Fig. 5d-e). Thus, it is possible that FGF23 activates promoter-enhancer long range interactions within these genes. To gain insight into the overall transcriptional regulatory state of cells treated with FGF23, the motif discovery algorithm (HOMER) was used to determine the enriched TF binding among open chromatin regions. This enrichment analysis showed high statistical significance for genome wide accessibility increases in the basic leucine Zipper domain (bZIP) factors including FOS, JUNB, and AP1, as well as an enhanced enrichment of EGR1 motifs (Supplementary Fig. 8b).

To determine the correlation of the epigenetic landscape and the global pattern of mRNA expression, we compared the derived ATACseq and RNAseq data via integration analyses. By plotting the chromatin accessibility for each gene *versus* its level of expression following FGF23 treatment, 349 genes were identified with significantly more open chromatin status (440 peaks) leading to higher gene expression levels after treatment (Fig. 5f), whereas 135 genes were downregulated with notably less DNA accessibility (153 peaks) (Fig. 5f). Among the genes that showed higher chromatin accessibility within a 10kb promoter region and associated with elevated gene expression were *FOSL1, NAB2, VDR, HBEGF,* and *NFKB1* (Fig. 5f). Functional analysis of differentially expressed genes at 4h after FGF23 treatment identified activated signaling pathways including FGFR1, NF-κB and MAPK (Fig. 5g). Among the genes associated with NF-κB signaling upregulated with FGF23 treatment were *BCL10, BCL2, CARD10, CXCL2, CXCL8, GADD45A, GADD45B, IL1R1, MAP3K14, PTGS2, RELB,* and *TNFRSF1A* (Fig. 5h). Specifically related to NF-κB signaling, a positive regulation of I-kappa B, involved in propagating the cellular response to inflammation, was predicted to be activated, supported by the increase of target genes such as *CTH, CCR7, CXXC5, HMOX1, MAP3K14, TRIML2, IL1A, BCL10, DDX21,* and *TNFRSF1A* (Fig. 5h).

To test the interrelationships between MAPK and NF-κB in the context of FGF23 signaling, a selective inhibitor of I-kappa B kinase phosphorylation^56^, BMS-345541, was used. Pretreatment of HEK293-mKL cells with BMS-345541 for 1h followed by FGF23 treatment for 4h resulted in sustained FGF23-dependent MAPK bioactivity as evidenced by the increase of ERK1/2, c-JUN, and c-FOS phosphorylation as well as enhancing the stabilization of c-FOS protein (Fig. 5i). Further, qPCR analyses showed that NF-κB inhibition increased FGF23-dependent *EGR1* and *FOS* mRNA expression (Fig. 5j). Taken together, these data support an intrinsic mechanism whereby FGF23-induced NF-κB activity acts as a negative regulator of FGF23/KL-driven MAPK signaling.

To test the hypothesis that FGF23 drives MAPK and NF-κB signaling in parallel but that NF-κB is inhibitory, we built upon the *in vitro* data and performed two *in vivo* experiments (Fig. 5k). First, wild type mice were treated with BMS-345541 or vehicle for 1h followed by FGF23 treatment for 1h. Our data showed that BMS-345541 administration promoted FGF23-induced renal *Fos* and *Egr1* mRNA expression. This approach also potentially enhanced 1,25D suppression by FGF23 through further decreasing Cyp27b1 mRNA expression when compared to mice treated with FGF23 alone (Fig. 5l). The blockade of NF-κB signaling also suppressed basal Cyp24a1 mRNA, potentially limiting FGF23 mediated increases. Secondly, a mouse model of X-linked hypophosphatemia (XLH), the *Hyp* mouse, known to closely mimic the human phenotype of this disease was used. This model manifests elevated FGF23, inappropriately normal 1,25D for the degree of hypophosphatemia, and reduced kidney sodium-phosphate transporter *Slc34a1* (Npt2a) expression^57^. *Hyp* mice treated with BMS-345541 for 3h had increased renal FOS and EGR1 mRNA versus controls (Fig. 5m) and normalized Cyp27b1 and Cyp24a1 alterations observed in this model. Further, investigation of Npt2a by immunofluorescence in kidney cortex supported that a chronic treatment of normal mice with BMS-345541 may be required to reach a Hyp mouse-level Npt2a decrease and activity (Supplementary Fig. 9). Collectively, these data support that NF-κB is a novel regulator of FGF23-dependent MAPK activity and kidney 1,25D metabolic enzymes.

### TNF-related family members as novel KL-dependent FGF23 gene targets

Elevated FGF23 production has been associated with pro-inflammatory states including during the chronic inflammation and hyperphosphatemia that are both associated with CKD. Although there have been several hypotheses proposed,^58,59^ the molecular mechanisms driving these FGF23-mediated crossover responses in kidney remain unclear. In this regard, we sought to isolate inflammatory biocomponents regulated by FGF23 bioactivity. Upon differential gene expression analysis within the kidney at the single cell level, we found a pronounced increase of TNF-associated genes following extended FGF23 delivery. These included elevated TNF receptor superfamily members in the PT, DCT, and CNT, which corresponded with the expression of KL. In response to FGF23, the maximal expression of the receptor member 12A (*Tnfrsf12a*) and 1A (*Tnfrsf1a*) was detected at 4h (Fig. 6a). In contrast, the renal expression of TNF ligands *Tnfα* and *Tnfsf12* was not induced by FGF23 in epithelial or immune cells, such as monocytes (Supplementary Fig. 8). The upregulation of TNF receptors *Tnfrsf1a* and *Tnfrsf12a* was restricted to PT-S1 and DCT, suggesting a potential KL-dependent activity and a coordinated role between renal epithelial and immune cells under inflammatory conditions in regulating FGF23 bioactivity and renal immune responses through TNFR signaling.

**Fig. 6:**
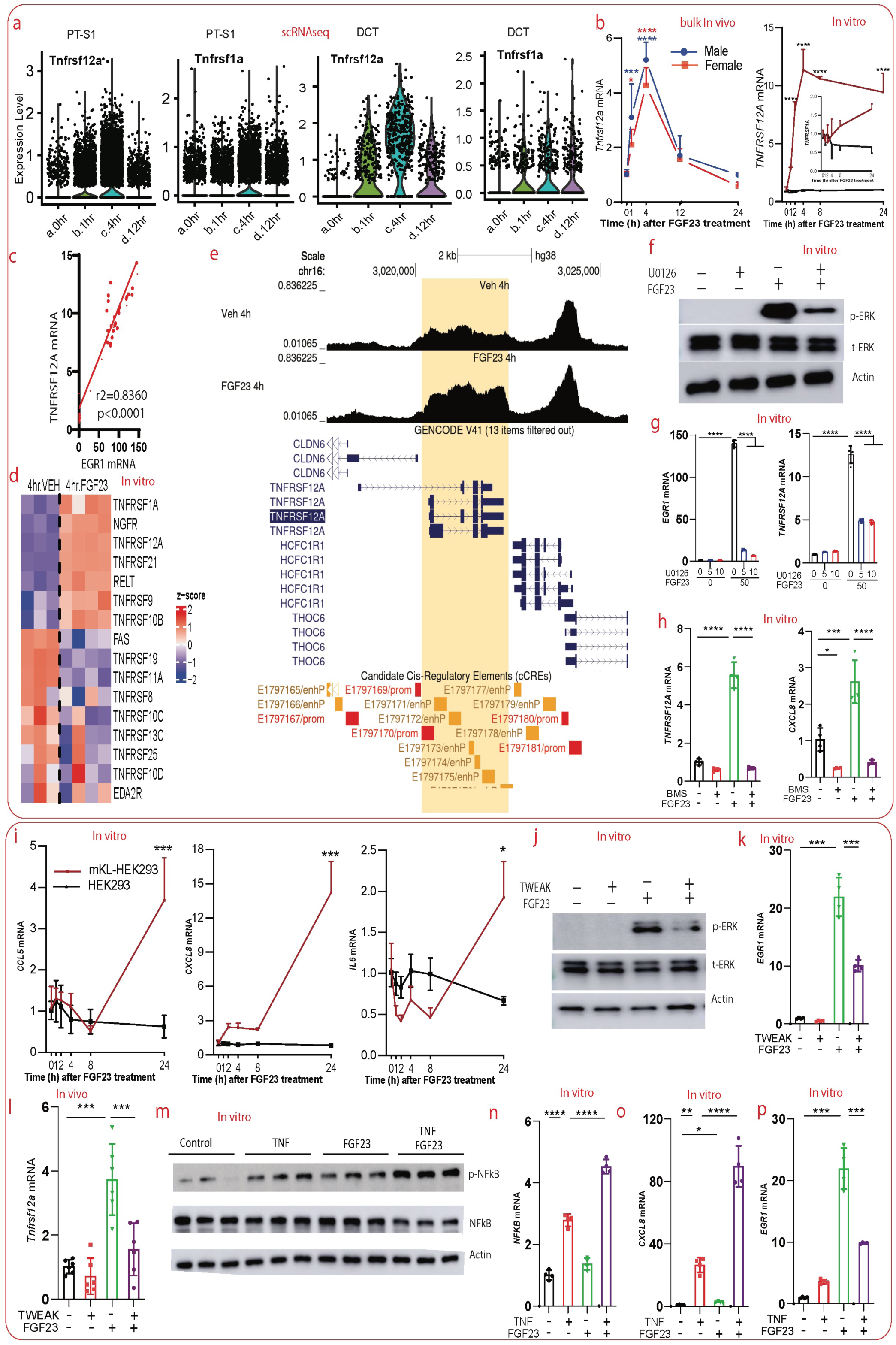
KL-dependent FGF23 bioactivity regulates Tnfrsf12a. **a.** Violin plots display the expression of *Tnfrsf12a* and *Tnfrsf1a* in PT-S1 and DCT. The kinetic curves show *Tnfrsf12a* mRNA expression in total kidney samples in male (blue) and female (red) mice treated with FGF23 for 0, 1, 4, 12, and 24h. **b.** In vitro analysis highlights the expression of *TNFRSF12A, EGR1*, and *TNFRSF1A* in HEK293 (black line) and HEK293-mKL (red line) cells treated with FGF23 for different times. **c.** Correlation analysis shows the positive interrelationship between *EGR1* and *TNFRSF12A.* **d.** HEK293-mKL cells were treated with FGF23 (50 ng/ml) for four hours and RNAseq was performed. The heatmap graphic displays differential gene expression of TNFRs genes that change with FGF23 treatment. **e.** HEK293-mKL cells were treated with FGF23 (50 ng/ml) for 4 hours and ATACseq was performed. Representative ATACseq peaks of cells treated with vehicle (top track) compared to cells treated with FGF23 (lower track) highlights the increase of chromatin accessibility across the TNFRSF12A gene body. **f.** HEK293-mKL cells were pretreated with the MEK inhibitor U0126 (5, or 10 μM) for 1h prior to FGF23 administration (50 ng/ml) for 10 minutes; p-ERK immunoblot. **g.** HEK293-mKL cells were pretreated with U0126 (5, or 10 μM) for 1h prior to FGF23 (50 ng/ml) for 4 hours followed by RNA extraction and assessment of *EGR1* and *TNFRSF12A* mRNAs by qPCR. **h.** HEK293-mKL cells were pretreated with BMS-34554 (10 μM) for 1h followed by FGF23 (50 ng/ml) for 4h then *TNFRSF12A* and *CXCL8* mRNA expression analysis. **i.** HEK293-mKL cells were treated with 50 ng/ml of FGF23 for 0, 1, 2, 4, 8, and 24h. The mRNA expression of *CCL5*, *CXCL8,* and *IL6* obtained from *in vitro* studies were measured by qPCR. The red kinetic curve represents the HEK293-mKL cell studies whereas the black kinetic curve shows the data from HEK293 cell line experiments. **j.** HEK293-mKL cells were treated with the ligand of TNFRSF12A (TNFSF12, TWEAK 100 ng/ml) for 1h followed by FGF23 treatment for 4h and pERK immunoblot. **k.** HEK293-mKL cells were pretreated with TWEAK (100 ng/ml) for 1h followed by FGF23 (50 ng/ml) for 16h. EGR1 expression was then tested by qPCR to assess FGF23 bioactivity. **l.** *In vivo* studies were performed by treating C57BL/6 mice with recombinant mouse TWEAK for 4 days at the rate of one injection per day. On the fourth day mice received FGF23 treatment for 4h and *Tnfrsf12a* was assessed by qPCR. **m-p.** HEK293-mKL cells were pretreated with TNF (100 ng/ml) for 16 h followed with FGF23 (50 ng/ml) for 4 h. pNF-κB, total NF-κB, and β-actin were evaluated by immunoblot (**m**). The mRNA expression of NFKB (**n**), CXCL8 (**o**), and EGR1 (**p**) were assessed by qPCR.

To test *in vitro* whether TNFRs are direct target genes of KL-dependent FGF23 bioactivity, parent HEK293 cells and HEK293-mKL cells were treated with FGF23 (50ng/ml) for 1, 2, 4, 8, and 24 hrs. In response to FGF23, *TNFRSF12A* (Fig. 6b) and *TNFRSF1A* (Fig. 6b-inset) were increased. In contrast to the robust signaling observed in HEK293-mKL cells, in the absence of KL in HEK293 cells no increase of *TNFRSF12A* or *TNFRSF1A* was observed (Fig. 6b). At 4h, TNFRSF12A mRNA was positively correlated with EGR1 expression, highlighting TNFRSF12A as a potential novel FGF23 target gene in KL-positive cells (Fig. 6c). By screening the TNFRs that were differentially expressed with FGF23 treatment in the RNAseq dataset, gene expression of *TNFRSF1A, NGFR, TNFRSF12A, TNFRSF21, RELT, TNFRSF9,* and *TNFRSF10B* were significantly increased (Fig. 6d). Among the gene ontology (GO) pathways predicted to be increased by FGF23 bioactivity based on integrated RNAseq and ATACseq datasets, were functions related to immune response-regulated signaling, control of NIK/NF-kB signaling, TNF signaling, TWEAK signaling, and TNFR1-induced NF-κB pathways. As highlighted in Fig. 5f, the chromatin accessibility at TNFRSF1A, TNFRSF12A, TNFRSF10B, TNFRSF1B, TNFRSF21, and NFKB promoter regions were increased with FGF23 treatment in KL-positive cells. Particularly striking, the chromatin accessibility at TNFRSF12A was increased along the entire gene body (Fig. 6e).

To assess whether the regulation of *TNFRSF12A* by FGF23 was MAPK-dependent and downstream of pERK signaling, HEK293-mKL cells were pretreated with the MEK inhibitor U0126 followed by FGF23. As assessed by immunoblot, pERK was inhibited in U0126-treated cells (Fig. 6f) and confirmed as associated with *EGR1* mRNA suppression (Fig. 6g). Importantly, MEK inhibition partially suppressed *TNFRSF12A* (Fig. 6g) expression in response to FGF23. These findings implicate the KL and MAPK axes in the regulation of TNF-associated pathways. Further, the inhibition of NF-κB using BMS-345541 suppressed FGF23-induced TNFRSF12A mRNA and downregulated CXCL8 in response to FGF23 (Fig. 6h). Interestingly, the known TNFRSF12A and NF-κB target genes *CXCL8, CCL5,* and *IL6* were upregulated in response to chronic FGF23 treatment (24h) (Fig. 6i), during which the MAPK target *EGR1* returned to baseline (Fig. 4e).

Kidney diseases such CKD and acute kidney injury (AKI) are associated with marked inflammatory state during which elevated production of pro-inflammatory cytokines including TNF family members such as TWEAK (TNFSF12) and TNFα can be observed^60,61^. Since FGF23 induced TNFRs (Fig. 6a-b), including the TWEAK receptor TNFRSF12A, we hypothesized the presence of a coordinated crosstalk between FGF23 and TNF signaling pathways to regulate renal epithelial KL-dependent FGF23 bioactivity. To isolate the impact of TNF ligand signaling on FGF23 bioactivity, HEK293-mKL cells were treated with TNFSF12 (TWEAK) followed by FGF23. Interestingly, the co-treatment showed that cells with initial activation of TWEAK signaling were more resistant to FGF23 bioactivity. Indeed, TWEAK-treated cells showed partial suppression of FGF23-induced ERK1/2 phosphorylation (Fig. 6j), as well as enhanced EGR1 mRNA (Fig. 6k). To test the effects of epithelial TWEAK signaling on renal FGF23 bioactivity, wild type mice were treated daily with TWEAK for 4 days followed by FGF23 delivery. In confirmation of the scRNAseq in kidney *in vivo* and RNAseq in HEK293-mKL cells, 4 hours after FGF23 treatment renal Tnfrsf12a mRNA expression was increased. However, pretreatment of mice with recombinant mouse TWEAK completely suppressed FGF23-mediated *Tnfrsf12a* increases (Fig. 6l). There were no marked effects on Npt2a protein following 4h FGF23 treatment (see Supplemental Fig. 9). To isolate whether TNFα may play similar role as TWEAK in the regulation of FGF23 bioactivity, HEK293-mKL cells were pretreated with TNFα for 1 hour followed by FGF23. Immunoblots demonstrated that cells treated with FGF23 had increased phosphorylation of NF-κB (p65; Fig. 6m) consistent with the predicted NF-κB signaling activation. Further, the co-treated cells showed an additive effect as there was further increased phosphorylation of NF-κB associated with elevated of NF-κB targets *NFKB* and *CXCL8,* and a parallel decrease of the MAPK-dependent gene *EGR1* (Fig. 6n-p). Thus, both TWEAK and TNFα *via* NF-κB may suppress FGF23-induced MAPK activity, which supports that impairment of FGF23 bioactivity occurs due to prevailing inflammatory signaling.

In sum, our findings herein identified key molecular events involved in renal FGF23 bioactivity at single cell level. We used a novel sex-specific demultiplexing analysis and found that KL-FGF23 interactions were sex independent, but KL expression drove cis functional responses in proximal and distal tubule cells, including overlapping and unique signaling events within these nephron segments. Our studies also identified novel functional segment-specific pathways and identified FGF23-mediated transcription factor regulon induction with effects on KL-positive cells. Finally, we identified crossover NF-κB and MAPK-dependent interactions that may potentiate TNFR signaling and restrict FGF23 cellular responses, potentially critical for mineral metabolism during disease states.

## Discussion

The cell-specific actions of FGF23 *via* its co-receptor KL are largely unknown, and to date have been solely based upon assumptions of segment-specific candidate gene expression. A primary goal herein was to leverage scRNAseq capabilities to facilitate the identification of FGF23-dependent targets that may impact kidney disease. Our “single-cell sex-hashing” study design provided the ability to determine sexual-dimorphic effects within the same experiment while avoiding the variability associated with technical processing. Using this approach, we were able to demultiplex male and female cell type across renal cells using X- and Y-chromosome specific genes in addition to PT sex-biased gene expression. Consistent with previous reports^23,62^, we identified the highest divergence of male- and female-biased gene expression in the PT-S3 region. Collectively, our data support these observations and demonstrate that the FGF23 responses elicited in males and females are similar. Our transcriptomic data at the single cell level demonstrated that the control of phosphate metabolism by FGF23 requires a temporally coordinated set of nephron cell-specific programs. To this end, MAPK signaling regulons were identified in PT-S1 and DCT segments associated with KL-dependent FGF23 bioactivity. In agreement with these findings, studies using genetic mouse models^63^ suggested that the effects of FGF23 on phosphate homeostasis could be mediated by the activation of the transcription factor Egr1, a known responder to MAPK induction. Using a temporal approach, we found FGF23 to rapidly induce gene sets associated with MAPK signaling prior to the regulation of transcripts associated with the control of 1,25D metabolizing enzymes Cyp24a1 and Cyp27b1 in PT-S1, but not in DCT/CNT. Further exploration of these associations through pairing scRNAseq with snATACseq identified cell type-specific chromatin accessibility for KL, as well as these critical FGF23-controlled enzymes.

The FGF23 co-receptor KL is required for high affinity FGF23 bioactivity^13^. We found that KL is transcribed at highest levels in the more homogeneous DCT/CNT compared to the levels observed in heterogenous PTS1-S3 segments which possess cells with varying KL mRNA expression, consistent with prior molecular mapping of the nephron by bulk analysis of isolated tubule sections^9,47^. Our findings suggest that PT segments, comprising the predominant kidney cell type, may have a more broad but lower KL expression, however whether this provides the ability to be more rapidly adaptive to minute-to-minute changes in FGF23 and phosphate handling remains to be determined. To test the importance of KL for FGF23 bioactivity, sub-setting analysis was employed based upon single cell KL mRNA expression levels. This was performed by testing epithelial cell clustering changes following FGF23 delivery, which reflected the overall gene transcription dynamics. We found that in response to FGF23, *KL^high^* cells exhibited a marked and parallel phenotypic change in gene expression, whereas *KL^low^* cells showed an unchanged transcriptional profile, and clustered more closely with untreated cells. Moreover, *KL^high^* cells at each time point following FGF23 delivery segregated more closely to each other due to similar gene regulatory control than at the original initiation point of being defined by nephron segment cell type. These results support that in response to acute FGF23 treatment in PT and DT, FGF23 induces common pathways but stimulates cell-specific functions. Further, potentially due to higher expression of KL in the DCT segments, FGF23 bioactivity may be sustained longer in this renal compartment *versus* PT-S1 as evidenced by a delayed ‘baseline-reset’ of key FGF23 target genes including *Egr1* and *Hbegf* (Fig. 3). Of note, the direct intracellular FGF23 signaling effects in KL-expressing cells occurred with a dynamic and common modification of specific gene regulons in the PT-S1 and DCT. Among the ‘early phase’ regulons driven by FGF23 in both PT-S1 and DCT were *Egr1*, *Fos*, and *Jun*, whereas ‘late phase’ regulons were defined by *Stat3* and *Nfκb* (Fig. 5a and Supplementary Data 1 and 2). In the broader sense, as opposed to neutralization of circulating FGF23, it has been therapeutically difficult to directly target FGF23-responsive pathways in kidney. Thus, these findings support the idea that to develop therapies which focus on FGF23-dependent mechanisms, different targeting strategies may have to be employed depending upon the temporal gene regulation pattern across nephron segments. Further studies may consider analyzing the impact of different FGFRs signaling on FGF23 bioactivity since a recent study showed that FGF23 also utilizes FGFR3c/4 as cognate receptor to form FGF23-FGFR3c-KL-heparin sulfate (HS) and FGF23-FGFR4-KL-HS quaternary signaling complexes,^64^ and differential FGFR expression along the nephron was observed from our scRNAseq dataset (Supplementary Fig. 2).

The analysis of the nephron at the single cell level also permitted reductions in the ‘noise’ associated with bulk RNAseq and array analyses from confounding non-FGF23 responsive cell types. Through pathway analysis, inflammatory responders, including TNFRSF12A, as described above, was identified *in vivo* and confirmed *in vitro* as a novel receptor controlled by FGF23. Consistent with parallel KL expression, TNFRSF12A localized primarily to PT (S1, S2, S3), DCT, and CNT. It is known that transient activation of TNFRSF12A signaling promotes protective tissue responses, playing a beneficial role in tissue repair following acute injury^65^. However, excessive and/or persistent TNFSF12/TNFRSF12A activation under pathological conditions promotes inflammation leading to tissue damage and degeneration^66^. Although FGF23 induced the expression of TNFRs including TNFRSF12A, the ligands for TNFRs were not induced in any kidney cell following FGF23 delivery, in agreement with previous work that showed that TNFSF12 is primarily upregulated in immune cells^67^. Thus, our findings support that FGF23 may directly control responses in KL-expressing cells, and through these actions potentially influence phenotypes in cells that are not directly FGF23-responsive. Our pathway analyses identified TNF-associated family members and NF-κB driven regulons as being enhanced with FGF23 delivery. To determine whether FGF23 inducible activity could be influenced by NF-κB signaling, NF-κB was inhibited *in vitro* and *in vivo* in the presence of FGF23. Indeed, NF-κB inhibition followed by FGF23 treatment completely abolished FGF23-induced TNFRSF12A expression as well as enhanced acute and prolonged MAPK mRNA responses, demonstrating that NF-κB is a novel negative regulator of kidney KL-dependent FGF23 bioactivity. Therefore, it is plausible that during CKD, chronic inflammation associated with increased NF-κB activity could inhibit FGF23-dependent mineral metabolism, as well as its protective functions on maintaining serum phosphate and 1,25D. KL has been shown to be downregulated in CKD models^68^, leading to the hypothesis that CKD is a state of FGF23 tissue resistance. Whether these alterations in MAPK/NF-κB signaling also contribute to dysfunctional FGF23 bioactivity prior to, and during, reductions in renal KL expression remains to be determined.

Although this work has marked strengths, including the identification of novel mechanisms of FGF23 actions, our studies are limited by the acute, transient aspects of FGF23 signaling tested herein. This approach was primarily undertaken to reduce any secondary effects due to marked systemic changes in mineral metabolism that occur with prolonged FGF23 delivery. Additional studies could consider testing the chronic effects of FGF23 signaling and test how the acute signaling mechanisms found herein are compensated long term. These mechanisms may potentially be important in diseases such as XLH and CKD where FGF23 is elevated often throughout patients’ lives. Further, our work confirmed known reductions in Npt2a protein expression with FGF23 delivery, but combined proteomic analysis of total kidney with our transcriptional datasets could reveal how the identified transcriptional changes detected in these studies are reflected at the translational levels in broad terms.

In summary, novel FGF23 targets with renal physiological and pathological functions were identified in this work at the single cell level *in vivo* and confirmed *in vitro.* Thus, these studies provide critical insight into the biology of FGF23 bioactivity, which drove transcriptional re-programming in KL-positive epithelial cells. This work also identified that FGF23 initiates common upstream signaling pathways in PT and DT segments, followed by segment-specific events leading to differential control of mineral ion handling. FGF23 also induced a programmed chain of ‘early’ and ‘late’ phase transcriptional events, which revealed novel TNF family member activation of NF-κB associated with altered KL-dependent FGF23 bioactivity (see model in Fig. 7). Collectively, our studies support novel functions of FGF23 that may provide key junction points for interventions into diseases of mineral metabolism.

**Fig. 7.**
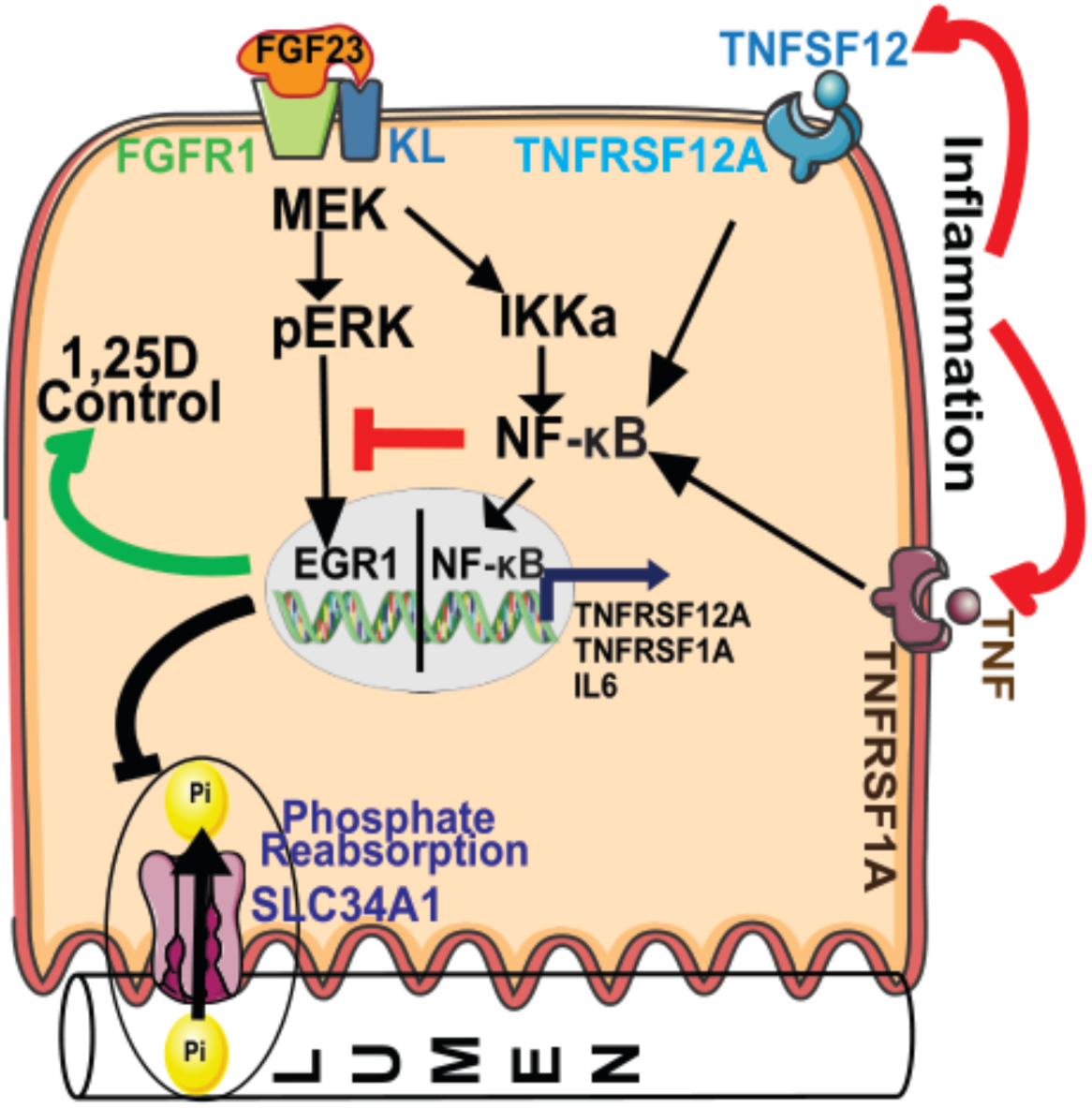
Proposed model. The graphic shows a proposed model of crosstalk between KL-dependent FGF23 bioactivity and inflammation-induced NF-κB signaling.

## Disclosure

KEW receives royalties for licensing FGF23 to Kyowa Hakko Kirin Co., Ltd; has current research funding from Calico Labs. KEW also owns equity interest in FGF Therapeutics. The other authors have nothing to declare.

## Acknowledgments

We thank Xuei Xiaoling, Patrick McGuire, and Hongyu Gao of the Center for Medical Genomics at the Indiana University School of Medicine for the assistance in performing single cell libraries preparation. We also thank the staff of IU Laboratory Animal Resource Center (LARC) for the care provided to animal throughout the studies. The authors would like to acknowledge NIH grants K99-DK129705 (RA), R01-DK112958, and R01-HL145528 (KEW), the David Weaver Professorship (KEW); R01-AI148282 (HT), K08-DK113223 (HT), the BX002901 Veterans Affairs Merit (HT). The content is solely the responsibility of the authors and does not necessarily represent the official views of the NIH or IUSM. We thank Drs. Nati Hernando and Carsten Wagner (University of Zürich, Zürich, Switzerland) for generously providing their anti-Npt2a antibody.

## Author’s Contributions

R.A. and K.E.W. designed and conceived the experiments; R.A., Y.G.M., P.N., M.L.N., and K.N.J. performed the in vivo and in vitro experiments; R.A. and D.J. conducted cell/nuclei preparation procedure for single cell biology experiments; T.H., J.M., S.L., and R.A. performed bioinformatics analysis and interpretated the data with advice from K.E.W., P.C.D., Y.L., and J.W; R. A. and K.E.W. drafted and wrote the manuscript. All the authors edited and approved the submitted content of the manuscript.

## Data availability

scRNAseq data is deposited in the NCBI’s Gene Expression Omnibus database (GEO GSE246385). The scATACseq analyzed in this paper were extracted from the GEO repository, accession number GSM6443123. The bulk RNAseq and ATACseq data presented are deposited in the GEO repository, accession number GSE254541.

## Supplementary Figures

**Supplementary Figure S1: Equal KL expression levels in male *versus* female renal cells.** Violin plots show KL expression in male and female kidney cells.

**Supplementary Figure S2: FGFRs expression in different kidney cells.** Violin plots show Fgfr1, Fgfr2, Fgfr3, and Fgfr4 mRNA expression in kidney cells.

**Supplementary Figure S3: snATACseq identified chromatin accessibility at VDRE binding elements within the KL genomic region of proximal tubule. a.** Unsupervised UMAP clustering identified renal cell clusters in the snATACseq dataset. Annotation was performed by label-transferring markers identified in the scRNAseq dataset. **b.** Differential chromatin accessibility is identified at the KL genomic region between exon 1 and exon 2 (chr5:150954700-150955500) highlighted in blue. The orange box pinpoints the absence of ATAC peaks between exon 1 and exon 2 in the DCT/LOH/CNT KL locus. **c.** Integrative Genomics Viewer (IGV) software was used to identify the corresponding binding regions. Within chr5:150954700-150955500, three publicly available ChIP-seq data (GSM3898510, GSM3898511, and GSM3898512) predict a VDRE binding domain specifically between chr5:150954800 and chr5:150955100. **c.** The dot plots represent gene activity of different kidney cell type markers.

**Supplementary Figure S4: DCT versus PT-S1 cell partition.** Monocle identified a single partition of DCT cells versus 6 different partitions in PT-S1 cells, highlighting the KL homogeneity vs the KL heterogeneity aspects in DCT versus PT-S1, respectively. **a.** Cells partitions using Monocle 3. **b.** FGF23 responses in DCT cells.

**Supplementary Figure S5: FGF23 activates common and specific pathways in PT and DT. a.** The heatmap highlights differentially expressed genes in PT-S1 and DCT. The red color represents upregulated genes (boxed in green) whereas the blue color highlights downregulated genes (boxed in blue). **b.** The violin plot shows the expression of Vdr in PT-S1. **c.** IPA analysis was used to identify the overlapping pathway network related to VDR/RXR activation (red box) in proximal tubule 1h after FGF23 treatment.

**Supplementary Figure S6: Kinetics of *in vivo* Npt2a protein regulation in response to FGF23 administration.** Images from kidney sections stained with Npt2a antibody (red) and DAPI (blue) revealed a pronounced intensity in kidney cortex (left). Enlargement of the kidney cortex showed a time-dependent decrease of Npt2a in response to FGF23.

**Supplementary Figure S7: SCENIC analysis of Egr1 and Fos transcription factors in DCT. a-b.** t-SNE plots illustrate *Egr1*, and *Fos* regulon activities in DCT cells. The histograms illustrate the AUCell score distribution for the regulon and the corresponding binding motifs. The mRNA expression of the transcription factor *Egr1* and *Fos* in DCT from SCENIC analysis and the scRNAseq dataset are shown respectively as a t-SNE feature plot with the orange dots displaying the mRNA expression levels. Violin plots show mRNA expression in the scRNAseq dataset.

**Supplementary Figure S8: ATAC-seq replicates and motifs enrichment. a.** Typical TSS enrichment plot shows that nucleosome-free fragments are enriched at the TSS. **b.** The motif discovery algorithm (HOMER) was used to determine the enriched TF binding among open chromatin regions. The dot plots show high statistical significance for genome wide accessibility increases in the basic leucine Zipper domain (bZIP) factors including FOS, JUNB, and AP1, and EGR1 motifs.

**Supplementary Figure S9: Effect of BMS and TWEAK treatments on Npt2a protein expression in the kidney.** C57BL/6 mice were treated with BMS, FGF23, or a combined BMS with FGF23, and frozen samples were sectioned and the slides stained with Npt2a antibody (red) and DAPI (blue). Klotho-KO, and Hyp mice were used as positive controls to confirm Npt2a detection. A quantitative approach (see Supplementary Methods) was used to measure the cortex fluorescence intensity over the medullary area (background).

**Supplementary Figure S10: Tnf ligands and receptor expression in monocytes.** Expression of *Tnf*, *Tnfrsf1a*, *Tnfrsf1b,* and *Tnfrsf12a* in monocytes in response to FGF23.

**Supplementary Table S1.** PCR Primers.

| Species | Genes | Primer References or Sequences | Vendor |
| --- | --- | --- | --- |
| Mouse | β-Actin | Mm02619580_g1 | ABI |
|  | Gapdh | Mm99999915_g1 | ABI |
|  | Kl | Mm00502002_m1 | ABI |
|  | Egr1 | Mm00656724_m1 | ABI |
|  | Cyp24a1 | Mm00487244_m1 | ABI |
|  | Cyp27b1 | Mm01165918_g1 | ABI |
|  | Slc34a1 (Npt2a) | Mm00441450_m1 | ABI |
|  | Tnfrsf12a | Mm07305008_m1 | ABI |
|  | Tnfsf12 | Mm02583406_s1 | ABI |
| Human | β-ACTIN | Hs1060665_g1 | ABI |
|  | EGR1 | Hs00152928_m1 | ABI |
|  | FOS | Hs00170630_m1 | ABI |
|  | BHLHE40 | Hs00186419_m1 | ABI |
|  | ATF4 | Hs00909569_g1 | ABI |
|  | DDIT3 | Hs01090850_m1 | ABI |
|  | TNFRSF12A | Hs00959047_g1 | ABI |
|  | TNFRSF1A | Hs01042313_m1 | ABI |
|  | CCL5 | Hs00982282_m1 | ABI |
|  | CXCL8 | Hs00174103_m1 | ABI |
|  | IL6 | Hs00174131_m1 | ABI |
|  | NFKB1 | Hs00765730_m1 | ABI |

## Supplementary Methods

### Immunofluorescence and imaging

Paraffin and frozen sections were stained using Npt2a primary antibody (1:1000; generously provided by Drs. Nati Hernando and Carsten Wagner) followed by a treatment with the polymer detection kit ImmPRESS® HRP Horse Anti-Rabbit IgG, Peroxidase (MP-7401; Vector Labs); subsequently the signal amplification kit TSA TMR reagent pack (Akoya Biosciences) was applied. Images were sequentially acquired in 2 separate channels using the Leica SP8 confocal microscope, collecting a single representative z-plane using a 20x NA 0.75 objective. Mosaics were stitched using Leica LAS X software to generate final images. A negative control without primary antibody was used to ensure the absence of nonspecific binding of secondary antibodies. Microscope settings were identical among imaging sessions for each specimen when imaging the same probes. Scale bar = 100 μm.

### Measuring cell fluorescence using ImageJ

We determine the level of cellular fluorescence from fluorescence microscopy images in ImageJ (http://rsbweb.nih.gov/ij/download.html). We select the cortex as region of interest and draw a polygon. Then we measure from the cortex fluorescence intensity. The background was measure in the medulla area. We then calculate the corrected total cell fluorescence (CTCF) using the formula CTCF = Integrated Density – (Area of selected cell X Mean fluorescence of background readings).

