## Supplemental file for "Dynamic Single Cell Transcriptomics Defines Kidney FGF23/KL Bioactivity and Novel Segment-Specific Inflammatory Targets"

Supplementary Fig. 1: Equal KL expression level in male versus female renal cells.

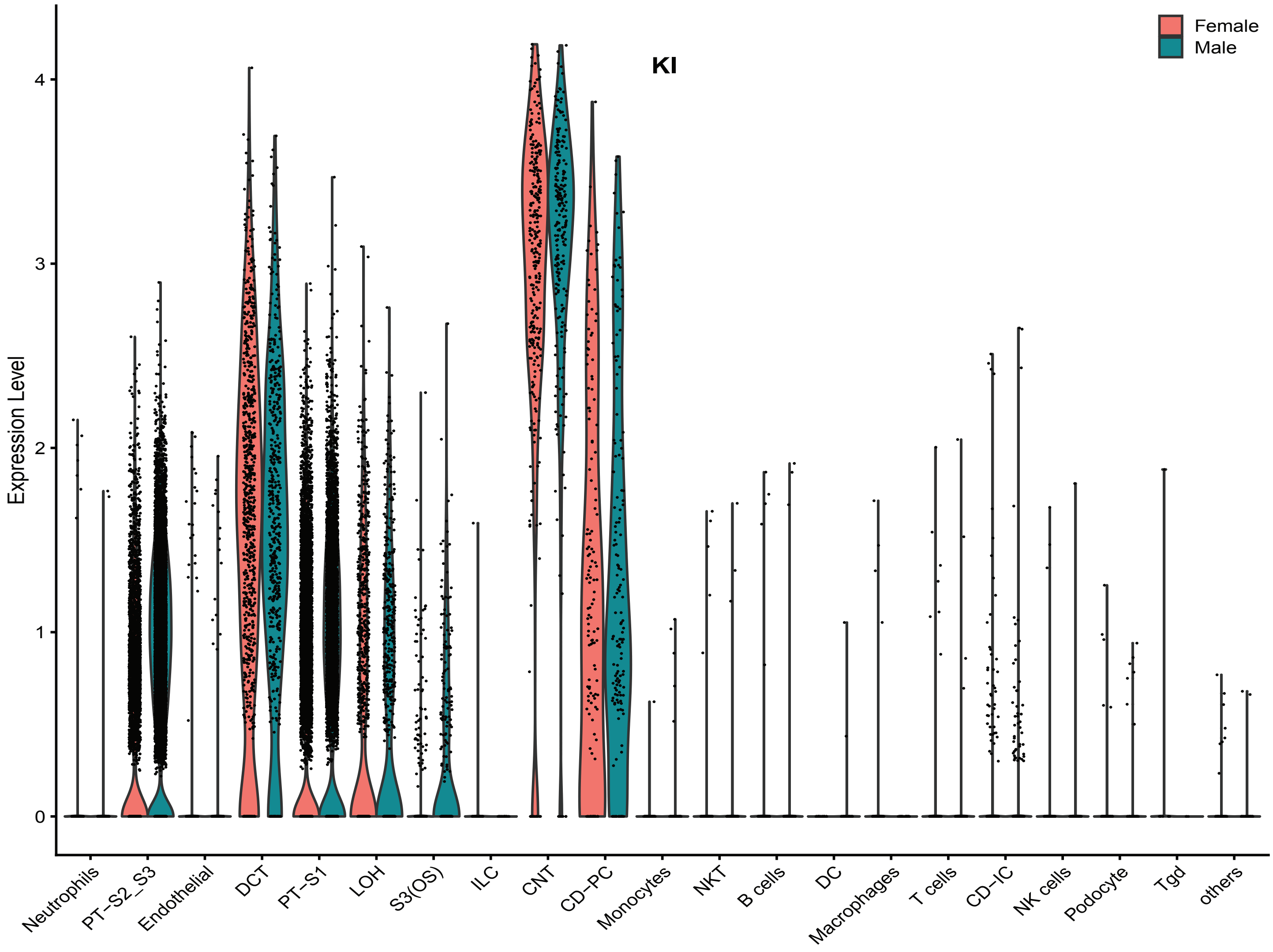

Supplementary Fig. 2: FGF receptors expression in renal cells.

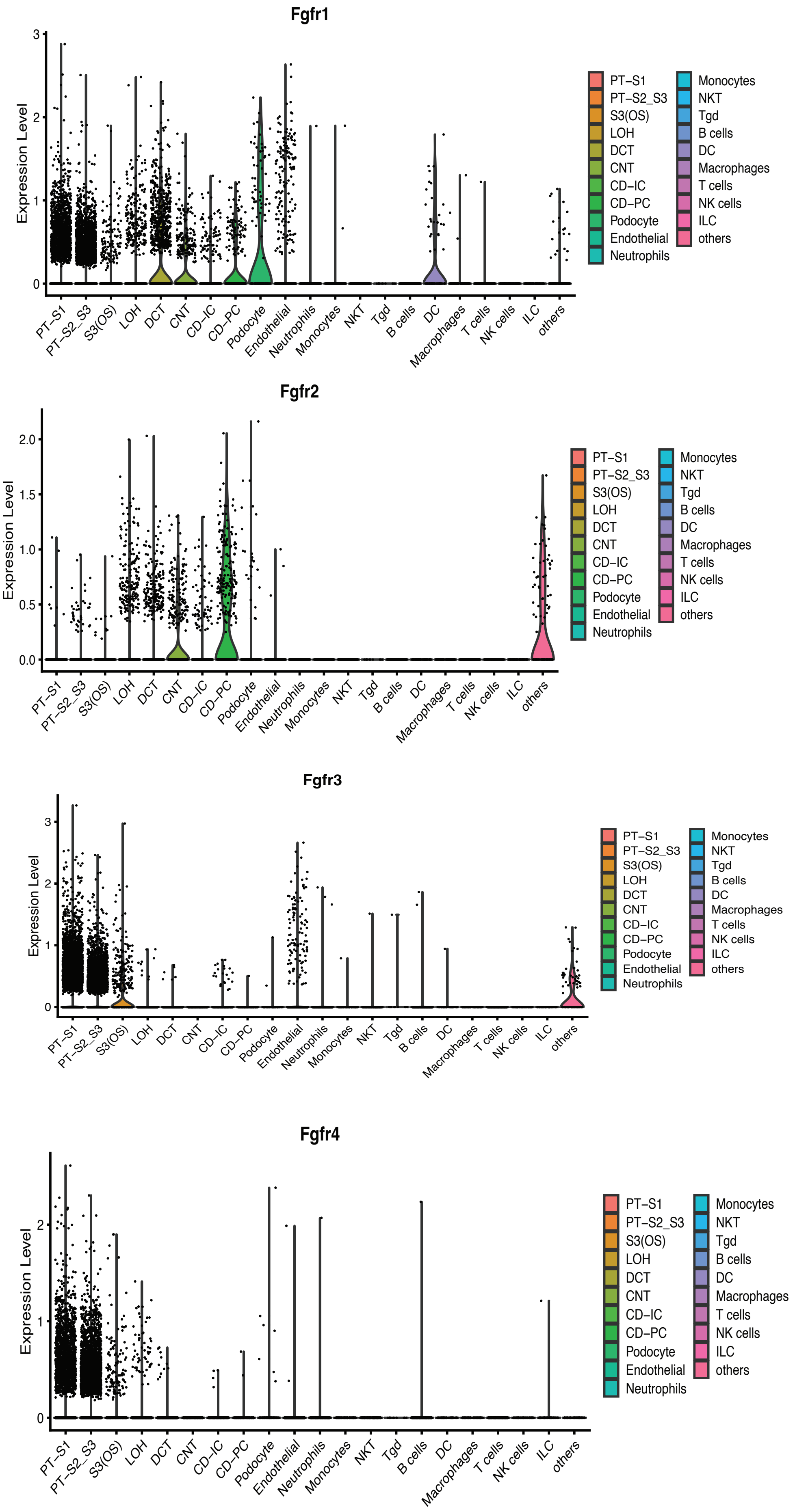

Supplementary Fig. 3: snATACseq identified a chromatin accessibility at VDRE binding elements within the Kl genomic region of proximal tubule

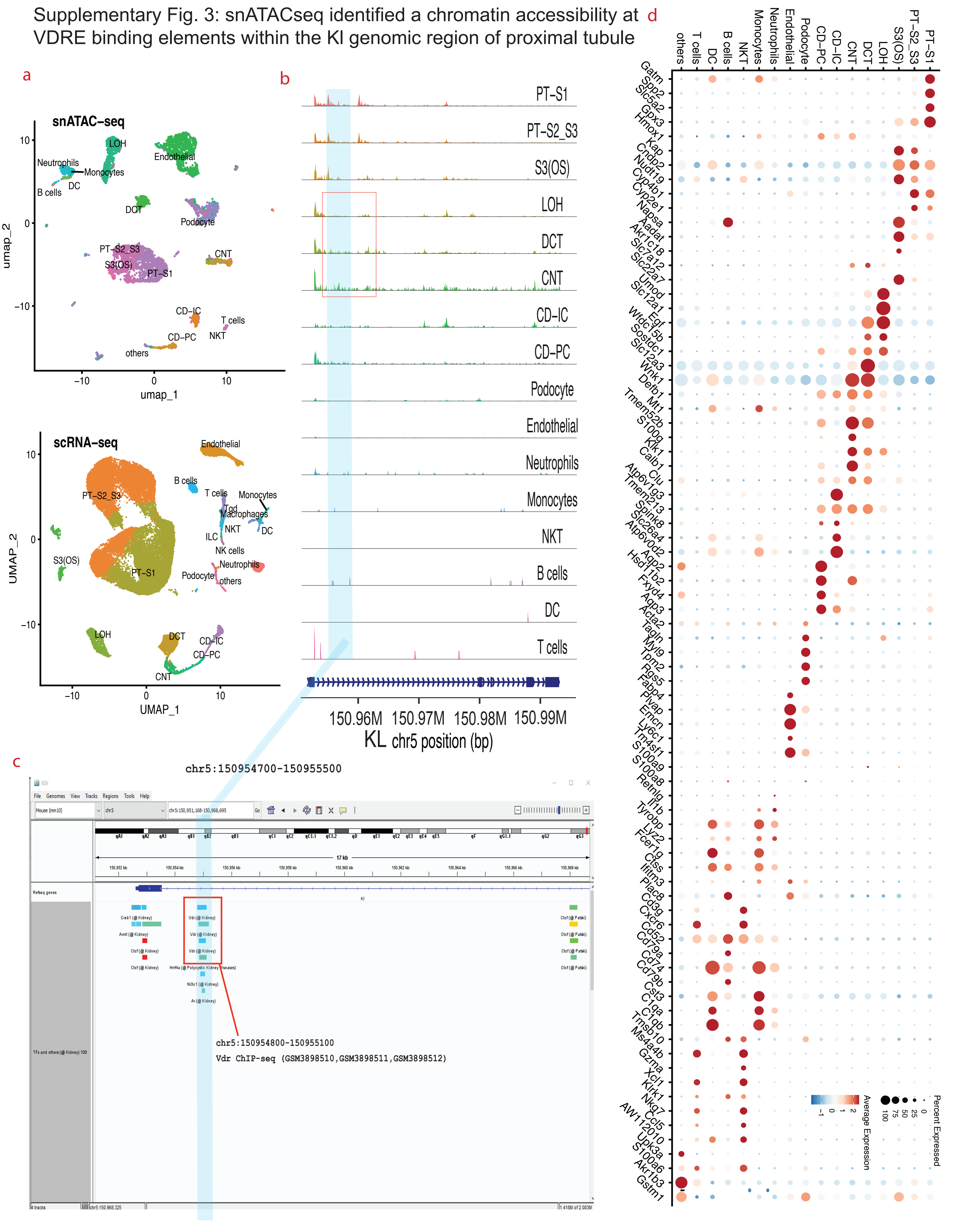

### Supplementary Fig. 4: DCT versus PT-S1 cell partition

a

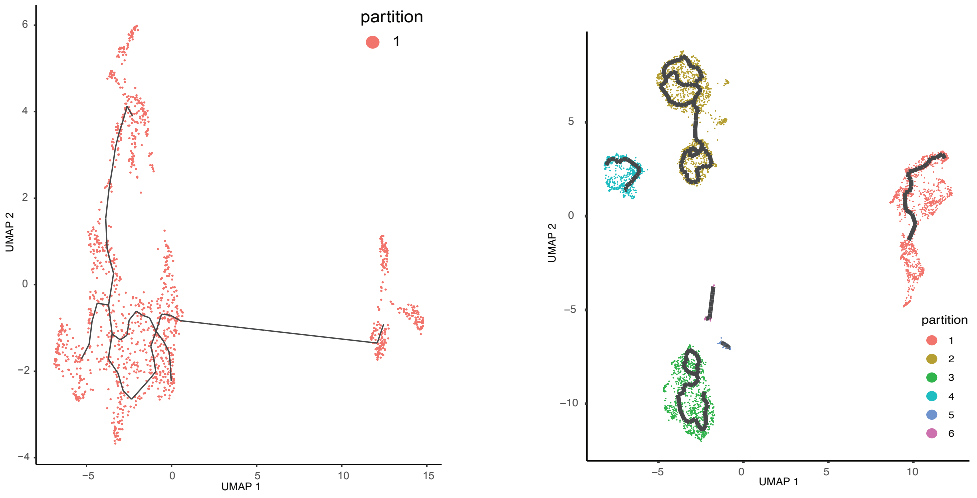


b

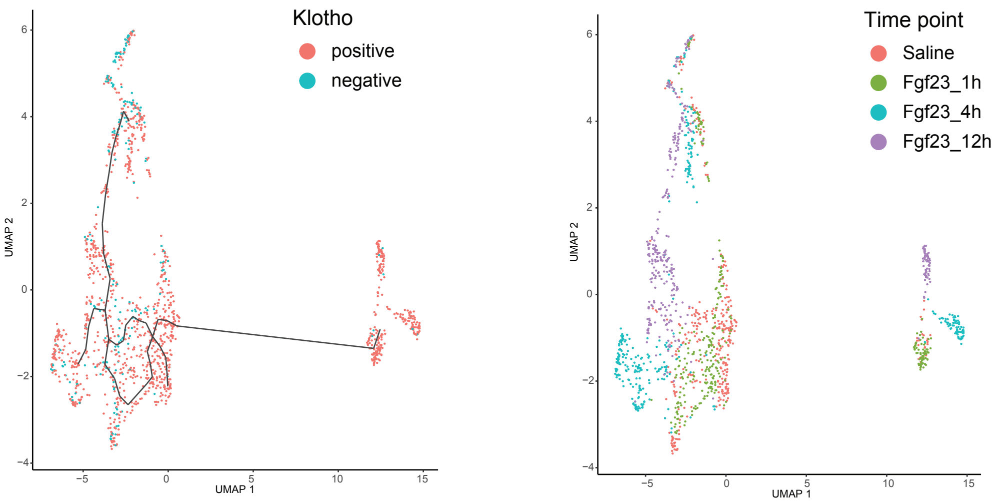

Supplementary Figure 5: FGF23 activates common and specific pathways in PT and DT.

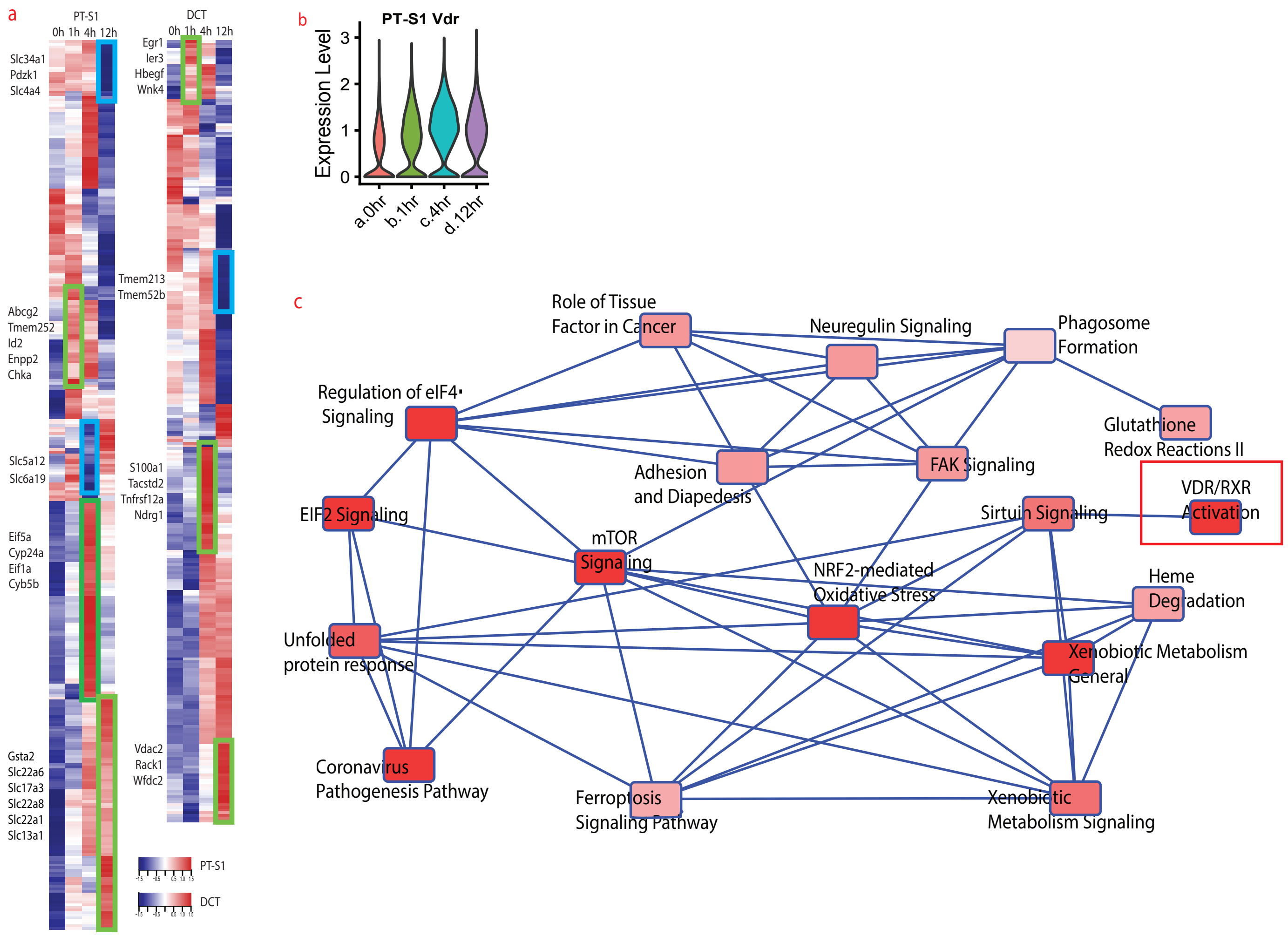

Supplementary Figure 6: Decrease of SLC34a1 protein in kidney in response to FGF23.

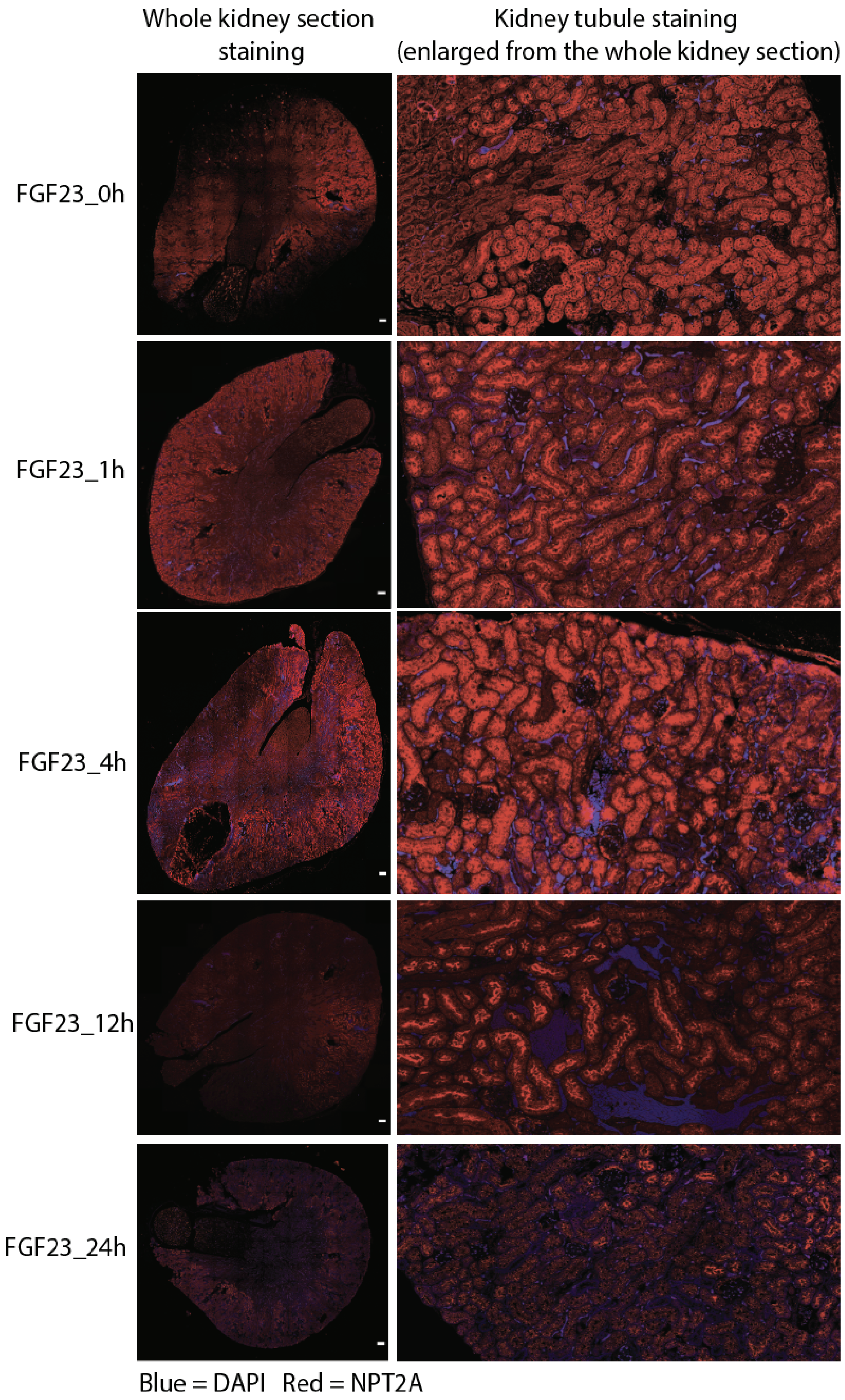

Supplementary Fig. 7: SCENIC analysis of Egr1 and Fos transcription factors in DCT

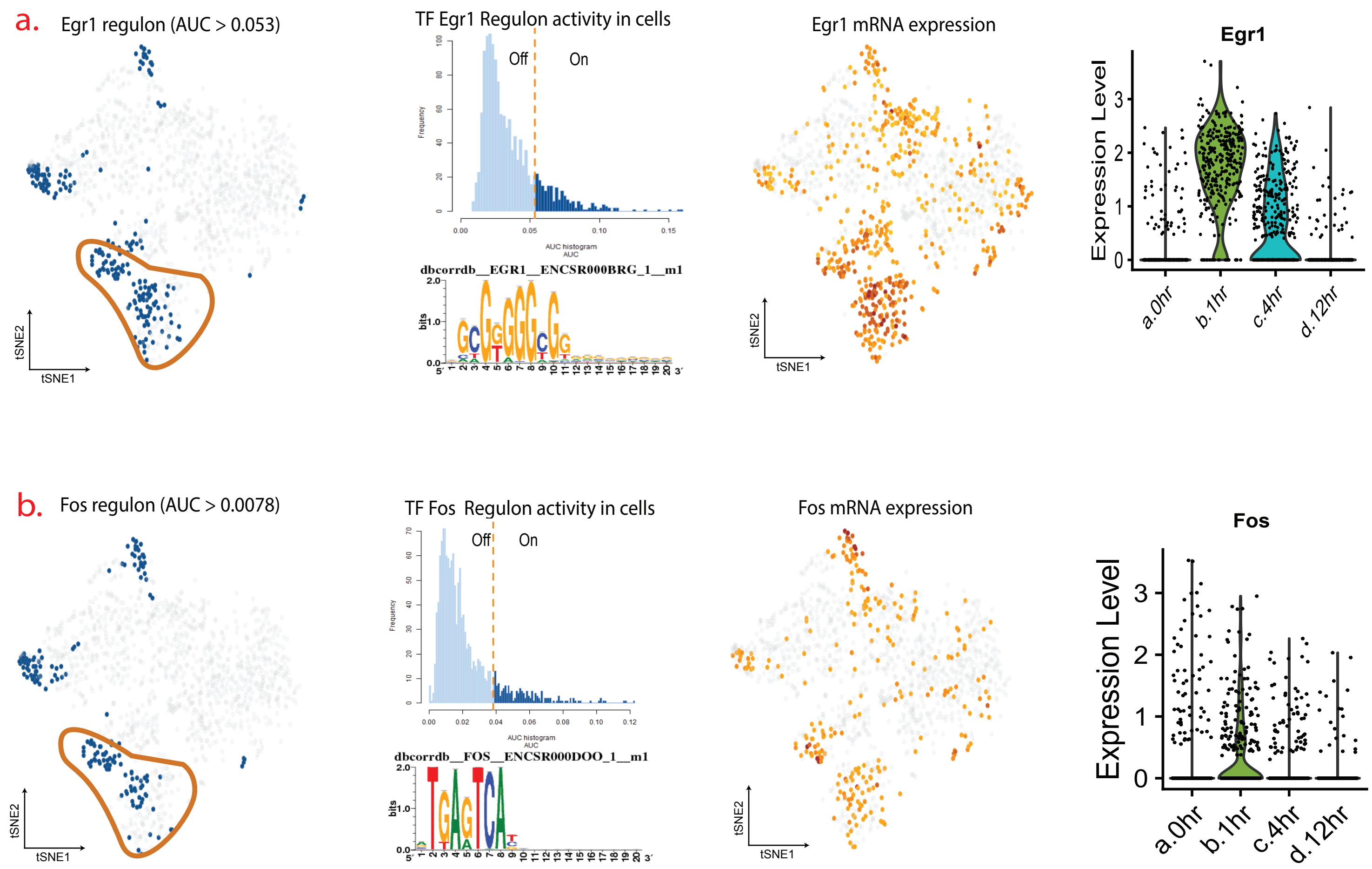

Supplementary Fig. 8: ATACseq replicates, and motifs enrichment

a.

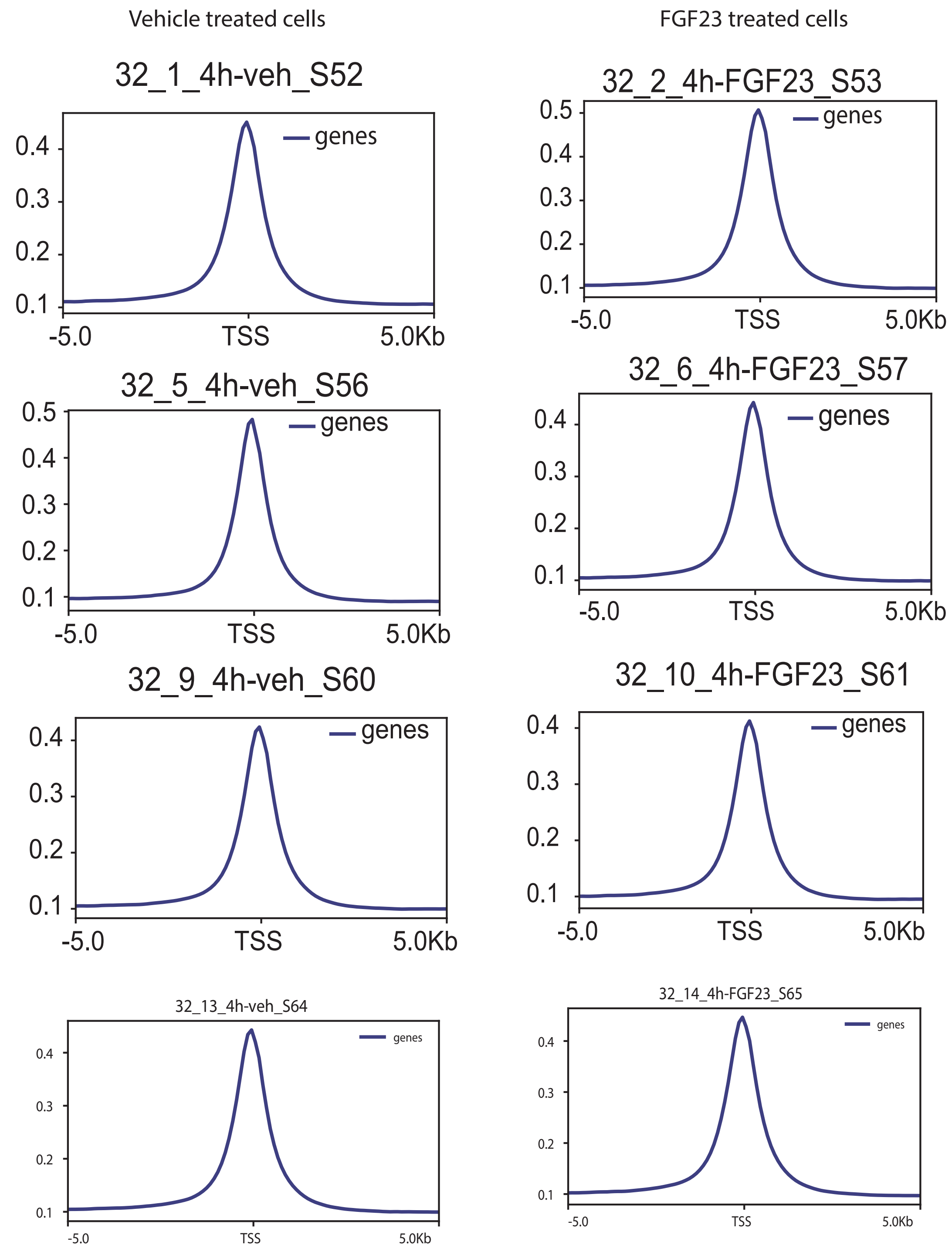


b.

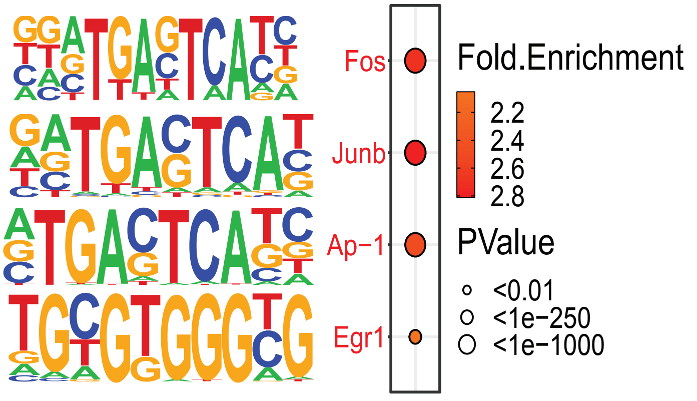

Supplementary Figure 9: Effect of BMS and TWEAK treatment on Npt2a protein expression in the kidney.

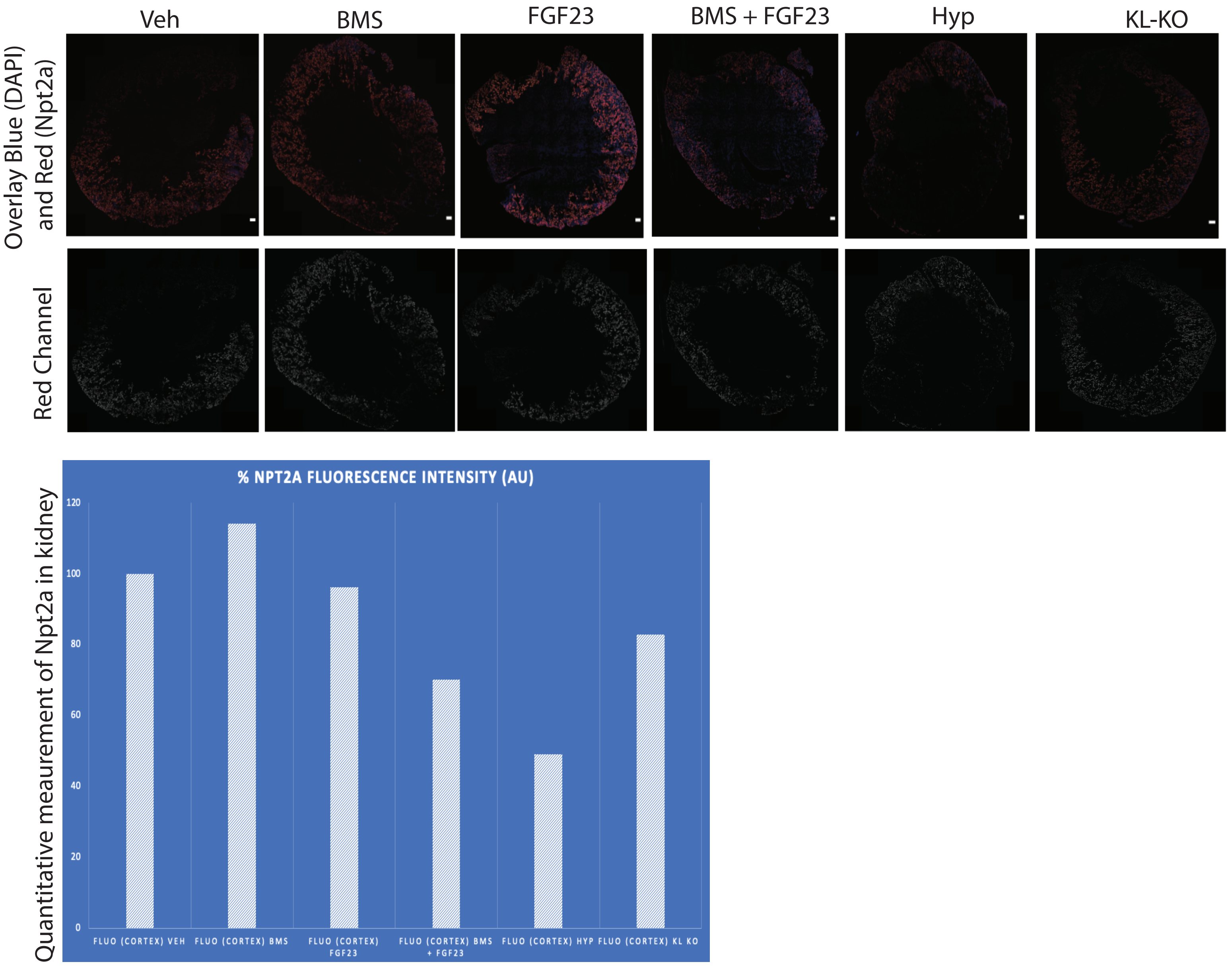


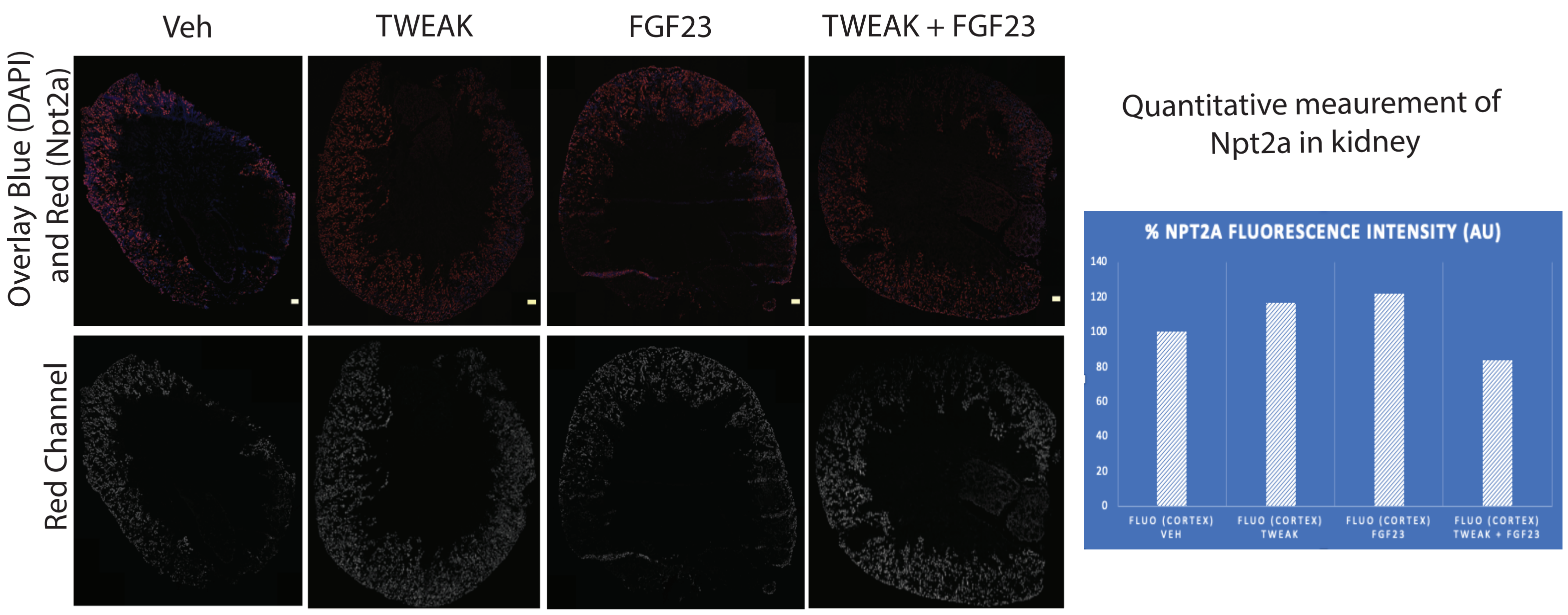

Supplementary Fig. 10: Tnf ligands and Tnf receptors expression in monocytes in response to FGF23.

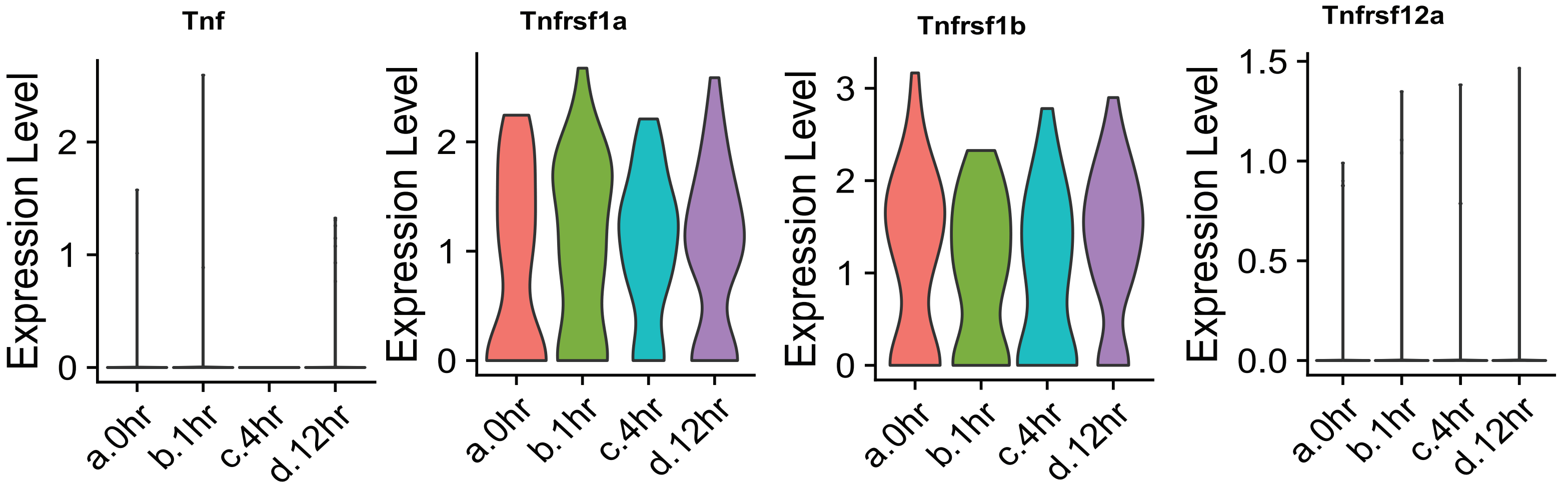

Supplementary Data 1 PT-S1 Regulon Activities

| Top | Regulon | FGF23 treatment (h) | RelativeActivity |
|---|---|---|---|
| 1 | Ets2_extended (11g) | a.0hr | 1.487800981 |
| 2 | Rcor1_extended (226g) | a.0hr | 1.472668993 |
| 3 | Srebf2 (207g) | a.0hr | 1.461742511 |
| 4 | Creb1_extended (1300g) | a.0hr | 1.455520602 |
| 5 | Zfp612_extended (171g) | a.0hr | 1.448047048 |
| 6 | Taf1 (1342g) | a.0hr | 1.441412471 |
| 7 | Yy1 (1990g) | a.0hr | 1.437843226 |
| 8 | Rbbp5_extended (70g) | a.0hr | 1.435070462 |
| 9 | Gtf2f1_extended (602g) | a.0hr | 1.434449654 |
| 10 | Maz_extended (3282g) | a.0hr | 1.431025214 |
| 11 | Atf1 (247g) | a.0hr | 1.419111193 |
| 12 | Esrra (10g) | a.0hr | 1.418584266 |
| 13 | Kdm5b_extended (381g) | a.0hr | 1.415094766 |
| 14 | Foxj3 (47g) | a.0hr | 1.414285062 |
| 15 | Phf8_extended (145g) | a.0hr | 1.412542042 |
| 16 | Foxj2_extended (13g) | a.0hr | 1.411975583 |
| 17 | Mga (13g) | a.0hr | 1.411039325 |
| 18 | Hcfc1 (1080g) | a.0hr | 1.407724781 |
| 19 | E2f4_extended (387g) | a.0hr | 1.404917722 |
| 20 | Mettl14 (12g) | a.0hr | 1.401079498 |

| Top | Regulon | FGF23 treatment (h) | RelativeActivity |
|---|---|---|---|
| 1 | Klf10_extended (18g) | b.1hr | 1.499745937 |
| 2 | Nr1h4_extended (21g) | b.1hr | 1.494648904 |
| 3 | Foxo3_extended (16g) | b.1hr | 1.480316361 |
| 4 | Bhlhe40 (13g) | b.1hr | 1.479987738 |
| 5 | Atf4_extended (13g) | b.1hr | 1.47525099 |
| 6 | Stat3 (28g) | b.1hr | 1.436895398 |
| 7 | Myc (53g) | b.1hr | 1.40472813 |
| 8 | Nr3c1 (4000g) | b.1hr | 1.392315984 |
| 9 | Junb (21g) | b.1hr | 1.36231885 |
| 10 | Egr1 (910g) | b.1hr | 1.155204573 |
| 11 | Jun_extended (290g) | b.1hr | 1.095180561 |
| 12 | E2f3_extended (13g) | b.1hr | 1.081220589 |
| 13 | Mafg (10g) | b.1hr | 1.073818977 |
| 14 | Fos (19g) | b.1hr | 1.031612956 |
| 15 | Jund_extended (2843g) | b.1hr | 0.989121835 |
| 16 | Hnf4a (512g) | b.1hr | 0.915724279 |
| 17 | Cebpb (54g) | b.1hr | 0.897251407 |
| 18 | Bhlhe41_extended (268g) | b.1hr | 0.876385676 |
| 19 | Fosb_extended (64g) | b.1hr | 0.859497343 |
| 20 | Hes1_extended (14g) | b.1hr | 0.849802566 |

| Top | Regulon | FGF23 treatment (h) | RelativeActivity |
|---|---|---|---|
| 1 | Irf8 (31g) | c.4hr | 1.470829081 |
| 2 | Nfe2l2_extended (44g) | c.4hr | 1.396801431 |
| 3 | Nfkb2 (71g) | c.4hr | 1.377994102 |
| 4 | Vdr_extended (31g) | c.4hr | 1.207020028 |
| 5 | Nfe2l1_extended (10g) | c.4hr | 1.151921385 |
| 6 | Hoxd9_extended (13g) | c.4hr | 1.082960458 |
| 7 | Xbp1 (81g) | c.4hr | 1.066829042 |
| 8 | Nfkb1 (51g) | c.4hr | 0.921104028 |
| 9 | Hes1_extended (14g) | c.4hr | 0.869258127 |
| 10 | Ppard_extended (14g) | c.4hr | 0.84437311 |
| 11 | Ppara_extended (33g) | c.4hr | 0.794336622 |
| 12 | Hnf4g_extended (15g) | c.4hr | 0.689003048 |
| 13 | Zmiz1 (21g) | c.4hr | 0.620434066 |
| 14 | Cebpb (54g) | c.4hr | 0.588313552 |
| 15 | Bhlhe41_extended (268g) | c.4hr | 0.378764269 |
| 16 | Mafg (10g) | c.4hr | 0.324089426 |
| 17 | Ar_extended (16g) | c.4hr | 0.242089987 |
| 18 | Cebpz_extended (2888g) | c.4hr | 0.157172808 |
| 19 | Nr3c1 (4000g) | c.4hr | 0.057379413 |
| 20 | Irf2 (12g) | c.4hr | 0.022561301 |

| Top | Regulon | FGF23 treatment (h) | RelativeActivity |
|---|---|---|---|
| 1 | Ar_extended (16g) | d.12hr | 1.255447242 |
| 2 | Hoxb7_extended (13g) | d.12hr | 1.253498256 |
| 3 | Zbtb11 (13g) | d.12hr | 1.242060535 |
| 4 | Nr2c1_extended (13g) | d.12hr | 1.129611242 |
| 5 | Elk1_extended (11g) | d.12hr | 1.042755882 |
| 6 | Foxo4 (109g) | d.12hr | 1.003914695 |
| 7 | Sap30 (245g) | d.12hr | 0.994357726 |
| 8 | Hnf4g_extended (15g) | d.12hr | 0.98823883 |
| 9 | Zfp143 (34g) | d.12hr | 0.943282986 |
| 10 | Polr3g_extended (19g) | d.12hr | 0.910917858 |
| 11 | Hoxc9_extended (16g) | d.12hr | 0.904547495 |
| 12 | Ppard_extended (14g) | d.12hr | 0.886519786 |
| 13 | Hes6_extended (22g) | d.12hr | 0.869740734 |
| 14 | Bach2_extended (17g) | d.12hr | 0.835553156 |
| 15 | Atf6 (71g) | d.12hr | 0.810280161 |
| 16 | Deaf1 (10g) | d.12hr | 0.749264938 |
| 17 | Gmeb2 (15g) | d.12hr | 0.740629699 |
| 18 | Polr3a (30g) | d.12hr | 0.725286202 |
| 19 | Hoxa5_extended (16g) | d.12hr | 0.618593155 |
| 20 | Hoxa10_extended (14g) | d.12hr | 0.601478485 |

Supplementary Data 2 DCT Regulon Activities

| Top | Regulon | CellType | RelativeActivity |
|---|---|---|---|
| 1 | Nfib (22g) | a.0hr | 1.485770738 |
| 2 | Klf16_extended (361g) | a.0hr | 1.48335759 |
| 3 | Batf3 (100g) | a.0hr | 1.455531435 |
| 4 | Tcf12 (29g) | a.0hr | 1.434001425 |
| 5 | Ppard_extended (33g) | a.0hr | 1.414264669 |
| 6 | Max_extended (1500g) | a.0hr | 1.384541863 |
| 7 | Klf15_extended (1661g) | a.0hr | 1.373734014 |
| 8 | Zfp110 (22g) | a.0hr | 1.372800344 |
| 9 | Zfp825_extended (18g) | a.0hr | 1.369893954 |
| 10 | Mxi1_extended (2012g) | a.0hr | 1.35857176 |
| 11 | Creb1_extended (1917g) | a.0hr | 1.347195144 |
| 12 | Hoxa5_extended (14g) | a.0hr | 1.333270048 |
| 13 | Mta3_extended (10g) | a.0hr | 1.332661555 |
| 14 | Nr2f2_extended (15g) | a.0hr | 1.314813196 |
| 15 | Rel (29g) | a.0hr | 1.310838471 |
| 16 | Bcl3_extended (17g) | a.0hr | 1.307736821 |
| 17 | Sin3a (575g) | a.0hr | 1.306064484 |
| 18 | Elf2 (1073g) | a.0hr | 1.304097416 |
| 19 | Pbx3_extended (145g) | a.0hr | 1.297158814 |
| 20 | Foxp1 (11g) | a.0hr | 1.292867254 |

| Top | Regulon | CellType | RelativeActivity |
|---|---|---|---|
| 1 | Crem (12g) | b.1hr | 1.49562859 |
| 2 | Klf10_extended (209g) | b.1hr | 1.486671101 |
| 3 | Nfil3 (24g) | b.1hr | 1.484852579 |
| 4 | Rela_extended (1148g) | b.1hr | 1.47814641 |
| 5 | Cebpg_extended (19g) | b.1hr | 1.46764252 |
| 6 | Ddit3_extended (21g) | b.1hr | 1.462798076 |
| 7 | Erf (701g) | b.1hr | 1.429448828 |
| 8 | E2f3 (60g) | b.1hr | 1.40346524 |
| 9 | Fosb_extended (199g) | b.1hr | 1.363311698 |
| 10 | Elf3_extended (576g) | b.1hr | 1.343251772 |
| 11 | Atf3 (189g) | b.1hr | 1.342140111 |
| 12 | Irf2_extended (22g) | b.1hr | 1.336966277 |
| 13 | Fos (71g) | b.1hr | 1.312538998 |
| 14 | Zmiz1_extended (6076g) | b.1hr | 1.298859061 |
| 15 | Hoxb6_extended (24g) | b.1hr | 1.279869707 |
| 16 | Egr1 (235g) | b.1hr | 1.277305218 |
| 17 | Foxo1 (277g) | b.1hr | 1.26550679 |
| 18 | Zfp513_extended (31g) | b.1hr | 1.222665822 |
| 19 | Klf9_extended (15g) | b.1hr | 1.217588269 |
| 20 | Smarcc2_extended (194g) | b.1hr | 1.206536836 |

| Top | Regulon | CellType | RelativeActivity |
|---|---|---|---|
| 1 | Vezf1_extended (14g) | c.4hr | 1.499334398 |
| 2 | Bmyc (19g) | c.4hr | 1.468464241 |
| 3 | Nr2f6 (10g) | c.4hr | 1.42244342 |
| 4 | Elk4 (1722g) | c.4hr | 1.377035514 |
| 5 | Xbp1_extended (38g) | c.4hr | 1.372721342 |
| 6 | Zfp217_extended (55g) | c.4hr | 1.300066725 |
| 7 | Stat3 (54g) | c.4hr | 1.297614671 |
| 8 | Srebf2 (290g) | c.4hr | 1.223336501 |
| 9 | Gtf2f1 (14g) | c.4hr | 1.160430763 |
| 10 | Nfe2l2_extended (14g) | c.4hr | 1.130512222 |
| 11 | Irf3 (1258g) | c.4hr | 1.126480365 |
| 12 | Hmgb1 (21g) | c.4hr | 1.092971744 |
| 13 | Cebpz_extended (1814g) | c.4hr | 1.011568229 |
| 14 | Jund (107g) | c.4hr | 0.967196054 |
| 15 | Kdm5b_extended (2079g) | c.4hr | 0.953902509 |
| 16 | Relb (90g) | c.4hr | 0.951811369 |
| 17 | Hoxb2_extended (20g) | c.4hr | 0.932838114 |
| 18 | Nfkb2 (127g) | c.4hr | 0.901210261 |
| 19 | Junb (131g) | c.4hr | 0.850588534 |
| 20 | Hes1_extended (15g) | c.4hr | 0.788709786 |

| Top | Regulon | CellType | RelativeActivity |
|---|---|---|---|
| 1 | Zfp143 (128g) | d.12hr | 1.486560951 |
| 2 | Maf (11g) | d.12hr | 1.473514038 |
| 3 | Foxk2 (16g) | d.12hr | 1.448542167 |
| 4 | Tead1 (52g) | d.12hr | 1.441192677 |
| 5 | Hnf1b (11g) | d.12hr | 1.429790272 |
| 6 | Zfp595 (21g) | d.12hr | 1.418713759 |
| 7 | Hdac2_extended (2768g) | d.12hr | 1.411533256 |
| 8 | Cic_extended (12g) | d.12hr | 1.403844208 |
| 9 | Tbp (23g) | d.12hr | 1.398262794 |
| 10 | Zscan12 (21g) | d.12hr | 1.391952839 |
| 11 | Mafg (17g) | d.12hr | 1.38898554 |
| 12 | E2f6_extended (1041g) | d.12hr | 1.38442944 |
| 13 | Xrcc4_extended (1013g) | d.12hr | 1.374091475 |
| 14 | Sp1_extended (14g) | d.12hr | 1.371322759 |
| 15 | Zfp846_extended (243g) | d.12hr | 1.364157238 |
| 16 | Tbx2 (11g) | d.12hr | 1.357856107 |
| 17 | Ctcf_extended (4122g) | d.12hr | 1.353071598 |
| 18 | Creb3_extended (20g) | d.12hr | 1.342948233 |
| 19 | Tia1 (11g) | d.12hr | 1.341126829 |
| 20 | Nrf1 (18g) | d.12hr | 1.340431761 |
